# Genetic causes and genomic consequences of breakdown of distyly in *Linum trigynum*

**DOI:** 10.1101/2023.11.16.567348

**Authors:** Juanita Gutiérrez-Valencia, Panagiotis-Ioannis Zervakis, Zoé Postel, Marco Fracassetti, Aleksandra Losvik, Sara Mehrabi, Ignas Bunikis, Lucile Soler, P. William Hughes, Aurélie Désamoré, Benjamin Laenen, Mohamed Abdelaziz, Olga Vinnere Pettersson, Juan Arroyo, Tanja Slotte

**Affiliations:** Department of Ecology, Environment and Plant Sciences, Science for Life Laboratory, Stockholm University, Stockholm, Sweden; Uppsala Genome Center, Department of Immunology, Genetics and Pathology, Uppsala University, Uppsala, Sweden; Department of Medical Biochemistry and Microbiology, Uppsala University, National Bioinformatics Infrastructure Sweden (NBIS), Science for Life Laboratory, Uppsala University, Uppsala, Sweden; Department of Genetics, University of Granada, Granada, Spain; Department of Plant Biology and Ecology, University of Seville, Seville, Spain

**Author notes:** Corresponding author: Tanja Slotte, Dept. of Ecology, Environment and Plant Sciences, Stockholm University, Stockholm SE-106 91 Stockholm, Sweden.

**Keywords:** homostyly, self-fertilization, distribution of fitness effects, genome assembly, plant mating system

## Abstract

Distyly is an iconic floral polymorphism governed by a supergene, which promotes efficient pollen transfer and outcrossing through reciprocal differences in the position of sexual organs in flowers, often coupled with heteromorphic self-incompatibility (SI). Distyly has evolved convergently in multiple flowering plant lineages, but has also broken down repeatedly, often resulting in homostylous, self-compatible populations with elevated rates of self-fertilization. Here, we aimed to study the genetic causes and genomic consequences of the shift to homostyly in *Linum trigynum*, which is closely related to distylous *Linum tenue.* Building on a high-quality genome assembly, we show that *L. trigynum* harbors a genomic region homologous to the dominant haplotype of the distyly supergene conferring long stamens and short styles in *L. tenue*, suggesting that loss of distyly first occurred in a short-styled individual. In contrast to homostylous *Primula* and *Fagopyrum*, *L. trigynum* harbors no fixed loss-of-function mutations in coding sequences of *S-*linked distyly candidate genes. Instead, floral gene expression analyses and controlled crosses suggest that mutations downregulating the *S-*linked *LtWDR-44* candidate gene for male SI and/or anther height could underlie homostyly and self-compatibility (SC) in *L. trigynum*. Population genomic analyses of 224 whole-genome sequences further demonstrate that *L. trigynum* is highly self-fertilizing, exhibits significantly lower genetic diversity genome-wide, and is experiencing relaxed purifying selection and less frequent positive selection on nonsynonymous mutations relative to *L. tenue*. Our analyses shed light on the loss of distyly in *L. trigynum*, and advance our understanding of a common evolutionary transition in flowering plants.

## Introduction

Flowering plants exhibit a remarkable diversity of floral and reproductive structures, in large part due to selection for increased pollination and reproductive efficiency (reviewed in Schiestl and Johnson 2013; Simón-Porcar et al. 2024). Understanding adaptation to different pollination modes is important, because this process can contribute to speciation, and is likely to underlie repeated convergent evolution of floral traits in flowering plants (reviewed in Barrett 2010, Ashworth et al. 2015). Evolutionary biologists have therefore long been fascinated by floral adaptations to pollination modes (Darwin 1876, 1877).

One such adaptation is distyly, a floral polymorphism that promotes efficient pollen transfer and outcrossing via insect pollinators (Darwin 1877, Lloyd and Webb 1992, Simón-Porcar et al. 2024, reviewed by Barrett 2019). Natural populations of distylous species are polymorphic for floral morph, such that individuals have one of two types of flowers that differ reciprocally in the positions of male and female sexual organs (anthers and stigmas, respectively) within the flower. Pin individuals have long styles (L-morph) and short stamens with stigmas at a high position and anthers at a low position in the flower, whereas thrum individuals have the reciprocal arrangement with short styles (S-morph) and long stamens (Fig. 1). In most distylous species, these morphological differences are coupled with a heteromorphic self-incompatibility (SI) system which allows inbreeding avoidance (Charlesworth and Charlesworth 1979b) and reinforces disassortative mating by preventing self- and intra-morph fertilization.

**Fig. 1.**
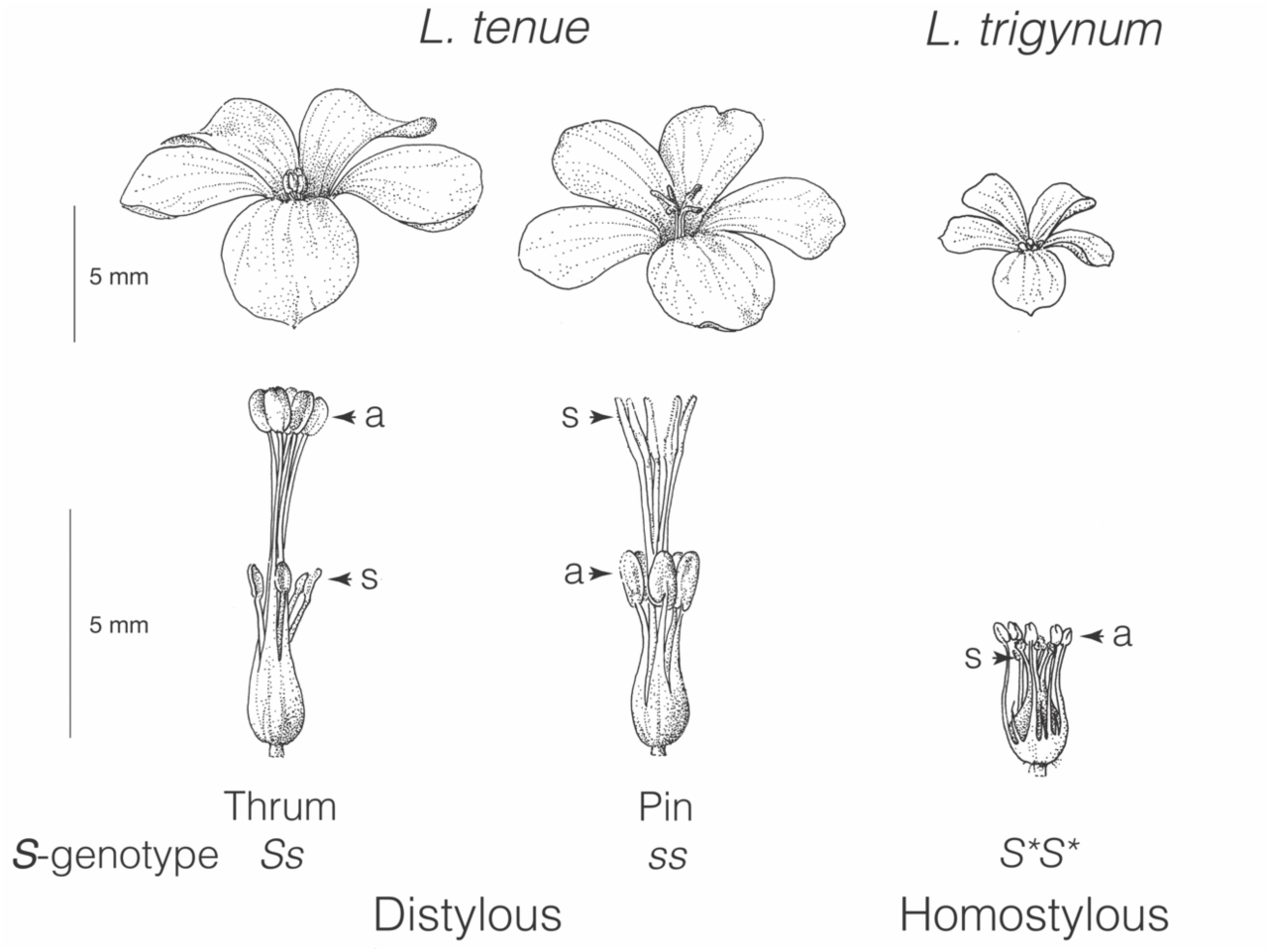
Flower morphs of the distylous *L. tenue* and the homostylous *L. trigynum*. The expected genotypes of thrum (left), pin (middle) and homostylous (right) flowers are indicated. In thrum flowers, pollen-producing anthers (indicated by a) are positioned above the short styles with receptive stigmas (indicated by s) in a low position in the flower, while pin individuals show the opposite reciprocal arrangement. Homostylous flowers show reduced herkogamy (i.e, anthers and stigmas in closer proximity). *S*= dominant haplotype, *s*=recessive haplotype, and *S\**= haplotype homologous to the *L. tenue S* allele. The position of stigmas and anthers are indicated in each morph. Illustrations by Alison Cutts.

Despite its multi-trait nature, distyly is inherited as a single Mendelian locus, called the distyly supergene or the *S-*locus, which governs both morphological differences and heteromorphic SI. Early work suggested that the distyly supergene typically harbors two alleles: one dominant exclusive to thrums and one recessive for which pins are homozygous (Bateson and Gregory 1905, Laibach 1923) (Fig. 1). It is only recently that distyly supergenes have begun to be sequenced and characterized in detail (reviewed in Gutiérrez-Valencia et al. 2021). So far, characterization of independently evolved distyly supergenes in six systems (*Primula*: Li et al. 2016, *Turnera*: Shore et al. 2019; *Linum*: Gutiérrez-Valencia et al. 2022a, *Fagopyrum*: Yasui et al. 2012, Fawcett et al. 2023, *Gelsemium*: Zhao et al. 2023; *Nymphoides indica*: Yang et al. 2023) suggest a remarkable degree of convergence in the genetic architecture of distyly supergenes. Common features of independently evolved distyly supergenes include a genetic architecture featuring length differences between the dominant and recessive supergene haplotypes, resulting in hemizygosity in thrum individuals and thrum-specific expression of *S-*linked genes (Gutiérrez-Valencia et al. 2022a). In addition, elevated repeat content is frequently found at distyly supergenes (Gutiérrez-Valencia et al. 2022a).

Style length polymorphisms, including distyly, have evolved independently multiple times in independent flowering plant lineages (Naiki 2012, Simón-Porcar et al. 2024). However, distyly has also broken down frequently, often resulting in homostylous species that are monomorphic and self-compatible (SC) with anthers and stigma at the same level (non-herkogamous; reviewed by Ganders 1979; note that here we restrict the use of the term homostylous to derived, monomorphic populations or species with anthers and stigmas at the same height, as in Darwin [1877]). Given their capacity for self-fertilization, facilitated by the joint occurrence of reduced herkogamy and SC, homostylous individuals (homostyles) can be favored whenever insect-mediated pollination and fertilization becomes unreliable, due to selection for reproductive assurance (e.g, Piper et al. 1986; Yuan et al. 2017). Genomic characterization of transitions from distyly to homostyly may therefore allow the identification of genetic changes underlying this mating system shift, as well as its population genomic consequences.

Because early models of the distyly supergene posited that thrum plants were heterozygous at the distyly supergene (Ernst 1936), it was frequently hypothesized that rare recombination events between dominant and recessive *S*-haplotypes caused homostyly (Dowrick 1956, Charlesworth and Charlesworth 1979a, reviewed by Ganders 1979). This idea has been revisited after the realization that thrums are predominantly hemizygous rather than heterozygous at the distyly *S*-locus in several distylous systems (Li et al. 2016, Shore et al. 2019, Gutiérrez-Valencia et al. 2022a, Fawcett et al. 2023). Since hemizygosity precludes the possibility of recombination between the dominant and recessive haplotypes, other genetic causes of distyly breakdown should be considered. First, genetic modifiers not linked to the distyly supergene could act by reducing floral herkogamy (Ganders 1979, Mather and De Winton 1941). Second, homostyly could evolve as a consequence of loss-of-function mutations at *S*-linked genes (as suggested by Ernst 1936). Such mutations could result in either long homostyles, with both anthers and stigma in a high position in the flower, or short homostyles, with their sexual organs in a low position in the flower.

Theory predicts that long homostyles, which exhibit functional dominant male SI in combination with recessive style length and female SI function should be favored during the establishment of homostyly, as they produce only homostylous offspring when pollinating compatible pin plants, in contrast to short homostyles which produce 50% homostylous offspring in crosses to compatible thrum plants (Dowrick 1956, Charlesworth and Charlesworth 1979a). As a consequence, long homostyles will spread faster than short homostyles at the initial stages of evolution of homostyly from distyly (although short homostyles can fix in the absence of long homostyles; Dowrick 1956, Charlesworth and Charlesworth 1979a).

In systems with hemizygous *S*-loci, it seems likely that homostyly would arise by mutation and not recombination. So far, results on the breakdown of distyly in *Primula* and *Fagopyrum* are in line with this prediction. In both systems, mutations affecting *S-*linked genes responsible for female SI function and short styles (i.e, thrum-exclusive *CYP734A50* in *Primula* encoding a brassinosteroid-inactivating enzyme, and *S-ELF3* in *Fagopyrum*) can readily lead to the formation of long-homostylous SC plants, because these *S*-linked genes jointly govern style length and female SI (Huu et al. 2016, Huu et al. 2022, Fawcett et al. 2023). Moreover, independently evolved homostyles in natural populations of *Primula vulgaris* harbor putative loss-of-function mutations in *CYP734A50* (Mora-Carrera et al. 2021). Mutations at *S*-linked genes, particularly in genes affecting style length and female SI, thus constitute a feasible pathway to the evolution of homostyly in natural populations, but it remains unknown if similar events have unfolded in lineages of distylous plants other than *Primula* and *Fagopyrum*.

Genomic studies hold the promise to elucidate potential genetic causes of loss of distyly and to quantify the impact of homostyly on outcrossing rates, as well as to characterize the consequences for patterns of polymorphism and the efficacy of selection. If the evolution of homostyly is associated with shifts to high selfing rates, we expect the transition to result in marked reductions in the effective population size (*N_e_*), exacerbated by linked selection due to reduced effective recombination rates in selfers, and potentially by founder events and bottlenecks associated with selfing (reviewed by Wright et al. 2013, Slotte 2014, Hartfield et al. 2017, Cutter 2019). In combination, these processes should result in reduced genetic diversity genome-wide in selfing homostylous species compared to their distylous relatives, which can also lead to more marked population structure and a decreased efficacy of selection, especially against weakly deleterious mutations but possibly also a reduced efficacy of positive selection. Although transitions from distyly to homostyly have occurred repeatedly in the history of flowering plants, the genomic consequences of this transition have so far primarily been studied in one system, *Primula* (e.g, Wang et al. 2021, Zhong et al. 2019).

*Linum* is a promising system for studying the evolution and breakdown of distyly (e.g. Armbruster et al. 2006, McDill et al. 2010, Gutiérrez-Valencia et al. 2022a, Innes et al. 2023), because it shows a remarkable diversity of stylar conditions, including several independent losses of distyly (Ruiz-Martín et al. 2018, Maguilla et al. 2021). In agreement with the expectation that selfers have improved colonization ability (Baker 1955, Stebbins 1957), phylogenetic analyses suggest that the evolution of homostyly is associated with the expansion of *Linum* outside its center of origin in the Western Palearctic (Maguilla et al. 2021). It remains largely unknown how homostyly has evolved in *Linum*, but we have recently shown that, similar to other distylous lineages, the distyly *S*-locus in *L. tenue* harbors a hemizygous region exclusively inherited by thrums (Gutiérrez-Valencia et al. 2022a). Importantly, this hemizygous region harbors a candidate gene for style length (*LtTSS1*), and the gene *LtWDR-44*, likely involved in anther elongation and/or pollen functioning (discussed in Gutiérrez-Valencia et al. 2022a). If mutations that impair *LtTSS1* function confer recessive style length and female SI function, we would expect such mutations to be favored during establishment of homostyly (Dowrick 1956, Charlesworth and Charlesworth1979a).

Here, we investigated the genetic causes and evolutionary genomic consequences of loss of distyly in *Linum trigynum*, an annual SC species distributed across the Mediterranean Basin and Western Asia (POWO 2024), in which homostyly is a derived state (Ruiz-Martín et al. 2018). For this purpose, we assembled and annotated a high-quality genome sequence of the homostylous *L. trigynum* using PacBio high-fidelity (HiFi) long reads and chromatin conformation capture (Hi-C) data. While a high-quality genome assembly is available for the closely related distylous *L. tenue* (Gutiérrez-Valencia et al. 2022a), no reference genomes of closely related homostylous *Linum* species were previously available. The new genomic resources for *L. trigynum* that we present here enable in-depth investigation of the evolution of homostyly and its genomic consequences.

To test hypotheses on the genetic basis of loss of distyly, we compared our *L. trigynum* genome assembly as well as two additional linked-read draft assemblies of *L. trigynum* to eight genome assemblies of the closely related distylous species *L. tenue* (Gutiérrez-Valencia et al. 2022a) to search for mutations at *S*-linked genes. We also made use of expression data to investigate patterns of differential expression at genes within the distyly supergene. To characterize the timing of the split between these species, we analyzed polymorphism data based on 224 whole genome sequences from eight natural populations each of *L. trigynum* and *L. tenue*. Finally, we conducted comparative population genomic analyses to test whether the shift to homostyly in *L. trigynum* was associated with reduced nucleotide diversity genome-wide, elevated inbreeding levels, increased population structure and a reduced efficacy of both purifying and positive selection, as predicted if the shift led to elevated selfing rates.

## Results

### A Chromosome-Scale Genome Assembly of Linum trigynum

To provide a genomic framework for studies of the evolution of homostyly in *L. trigynum*, we generated a high-quality *L. trigynum* genome assembly based on HiFi PacBio long reads (∼41x coverage) and Hi-C data from a single homostylous *L. trigynum* individual (see Materials and Methods and Table S1, Supplementary Information for sample information). The resulting assembly was highly complete (complete BUSCOs=96.6%; Table S2, Supplementary Information) and spanned 498 Mb (N50=47.03 Mb), divided in 10 pseudochromosomes and five scaffolds (less than 500 kb) with large-scale synteny to the genome assembly of distylous *L. tenue* (Gutiérrez-Valencia et al. 2022a) (Fig. 2a, Fig. S1, Supplementary Information). Our chromosome-scale assembly has the same number of pseudochromosomes as the haploid chromosome number in *L. trigynum* and *L. tenue* (*n=*10 for both diploid annual species; Valdés-Florido et al. 2023). Repetitive sequences constituted 56.49% of the assembly, and we annotated a total of 54,692 coding genes, similar to the number in *L. tenue* (Gutiérrez-Valencia et al. 2022a).

**Fig. 2.**
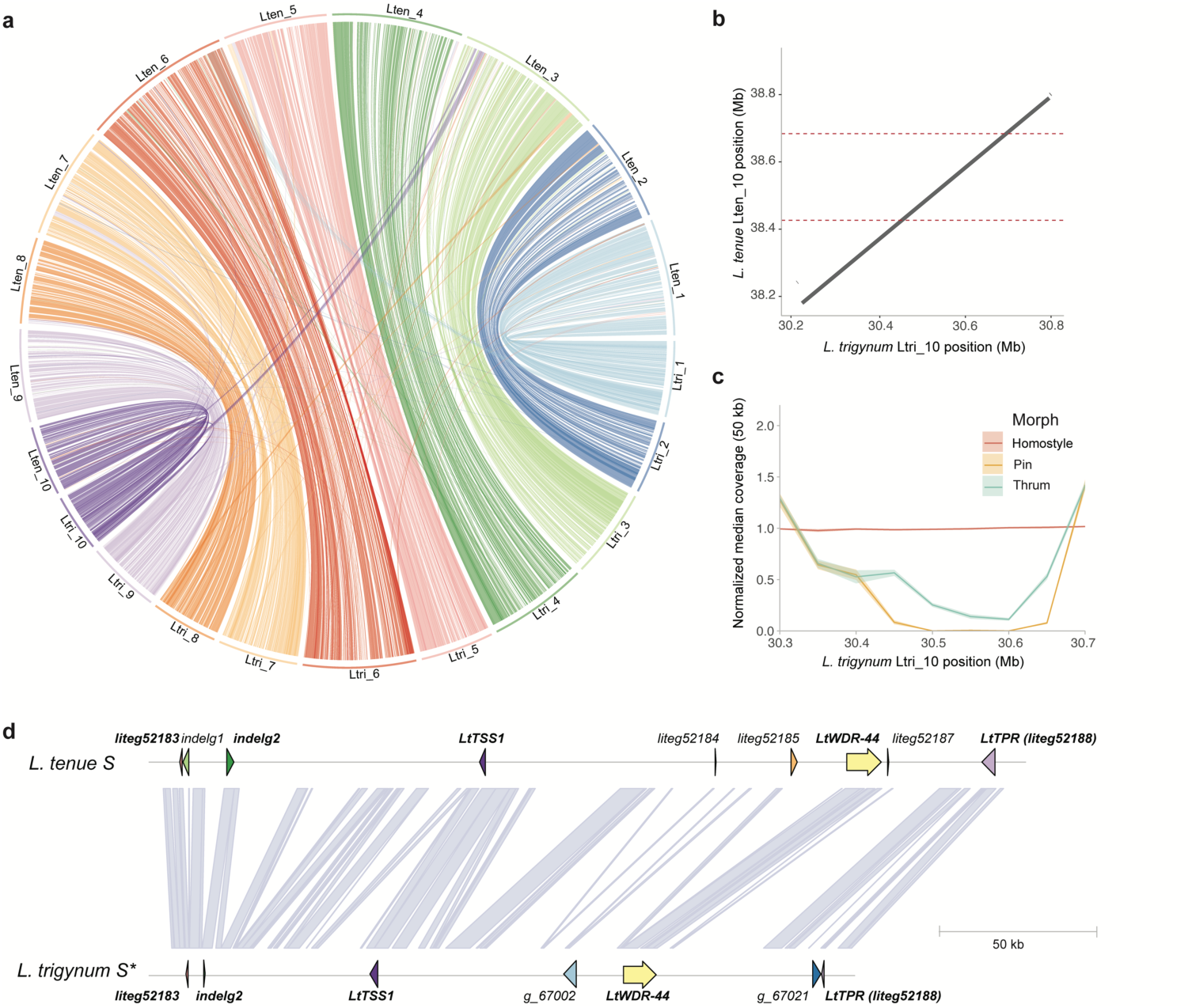
Broad-scale synteny between *L. trigynum* and *L. tenue* and comparative analyses of the *S-*locus region. **(a)** Circos plot showing broad-scale synteny between *L. tenue* (top) and *L. trigynum* (bottom). **(b)** Dotplot showing significant minimap2 alignments between the genomic region harboring the dominant *S-*haplotype of *L. tenue* and a region of *L. trigynum* chromosome 10. The limits of the *L. tenue S-*locus are indicated by dotted lines. **(c)** Plot of median and 95% confidence intervals of median coverage in 50 kb windows for *L. trigynum* (homostyle), *L. tenue* thrum and pin individuals, all mapped to our *L. trigynum* assembly. **(d)** Comparison of gene content at the *S-*haplotype of *L. tenue* (including genes *liteg52183* and *liteg52188* not within the hemizygous region of the *S-*locus; entire length of *S-*locus ∼261 kb) and corresponding region (here named *S\**; length ∼208 kb) in *L. trigynum*. Names of genes shared between the *L. tenue S-*haplotype and the *L. trigynum S\**-haplotype are written in bold text. Purple lines show orthologous regions between the two haplotypes.

We supplemented our PacBio genome assemblies of *L. trigynum* and *L. tenue* with two additional draft assemblies of *L. trigynum* based on 10x Chromium linked-read sequencing data (contig N50=4.82 Mb vs 4.47 Mb; assembly size=461.69 Mb vs 463.46 Mb), and seven additional *L. tenue* linked-read assemblies for thrum individuals which carry both the dominant and recessive haplotypes of the *S*-locus (Table S3, Supplementary Information).

### Analyses of the *S-*locus Region Elucidate the Origin of Homostyly

Whole-genome alignments of our high-quality *L. trigynum* and *L. tenue* assemblies showed that *L. trigynum* carries a region on chromosome 10 that is homologous to the longer, dominant allele of the *L. tenue* distyly *S*-locus (Fig. 2b). In contrast to *L. tenue*, which is polymorphic for a ∼260 kb indel at the *S-*locus (Gutiérrez-Valencia et al. 2022a), analyses of linked-read assemblies and genome coverage at this region based on short-read data from 104 *L. trigynum* individuals suggest that no such presence-absence variation is present in *L. trigynum*. Instead, all *L. trigynum* individuals harbor a haplotype similar to the longer *S*-haplotype that is dominant in *L. tenue* (Fig. 2c). This result suggests that events affecting the functioning of the supergene in thrum individuals led to the breakdown of distyly, and that the haplotype identified in *L. trigynum* is likely derived from the dominant *S-*haplotype.

We then investigated the gene content in the *L. trigynum* genome region homologous to the dominant *S-*haplotype of *L. tenue*. After curation of *S*-locus annotation (see Materials and Methods for details), we retained nine *S*-linked genes (Fig. 2d) in *L. tenue*. Out of these, five homologous *S-*linked genes were present in our high-quality long-read *L. trigynum* assembly (Fig. 2d). These included the thrum-specific gene *LtTSS1* which has pistil-specific expression and is likely to reduce cell length in thrum styles, and *LtWDR-44* potentially involved in controlling anther position and/or pollen incompatibility in *L. tenue* (discussed in Gutiérrez-Valencia et al. 2022a). In addition, *L. trigynum* harbored homologs of genes at the 5’ and 3’ ends of the *S-*locus region (*liteg52183* and *LtTPR*, respectively, Fig. 2d). The remaining genes, which are of unknown function (Gutiérrez-Valencia et al. 2022a), were not present in the *L. trigynum* genome assembly (Fig. 2d). Importantly, haplotype network analyses of *LtTSS1* and *LtWDR-44* indicate that variation at the *L. trigynum S-*locus stems from a single *S-*haplotype from its distylous ancestor (Fig. S2, Supplementary Information), and they further support the conclusion that loss of distyly first occurred in a short-styled individual, as suggested by the coverage-based analyses.

Next, we compared sequences of *L. tenue* and *L. trigynum* to identify candidate loss-of-function changes at the *S-*locus region affecting gene function in *L. trigynum*, and to quantify synonymous and nonsynonymous divergence between *L trigynum* and the dominant *S-*haplotype of *L. tenue.* We only identified putative loss-of-function changes in *L. trigynum* in the gene *LtTSS1* (Table 1). This gene harbors one exon, as demonstrated by our updated annotation, validated by PCR-assays of cDNA structure and similar to the gene structure of *TSS1* (*Thrum Style Specific 1*) in *L. grandiflorum* (Ushijima et al. 2012) (see Materials and Methods for details; Fig. S3 and Supplementary Methods; Supplementary Information). In *LtTSS1*, we identified a C-to-A mutation resulting in a premature stop codon in *L. trigynum,* that was located near the end of the predicted protein (eight codon positions prior to the regular stop codon), after the two conserved motifs previously identified in the *Linum grandiflorum* homolog *TSS1* (Ushijima et al. 2012). This C-to-A mutation was not fixed in *L. trigynum* but segregating at a frequency of 0.26 in our population data set. Taken together, the genic location and population frequency of this mutation suggests that it is unlikely to cause loss of distyly. Although all genes also harbored nonsynonymous mutations, there were no frameshift or other major-effect mutations in the coding sequence of the other analyzed genes, nor markedly accelerated rates of nonsynonymous substitution (Table 1). Finally, we used our repeat annotations to compare the transposable element (TE) content in the promoter region (2 kb upstream of the transcription start site) of *LtTSS1* and *LtWDR-44* in *L. tenue* and *L. trigynum*. We found no differences in the TE content in the promoter region of *LtTSS1* (Table S4; Supplementary information), but the TE content in the promoter region of *LtWDR-44* differed between species. *L. trigynum* harbored LTR/Ty3-like long terminal repeat (LTR) retrotransposon insertions in the *LtWDR-44* promoter that were not detected in *L. tenue*, and the type of DNA transposons detected also differed between the two species (Table S4; Supplementary Information).

**Table 1.**
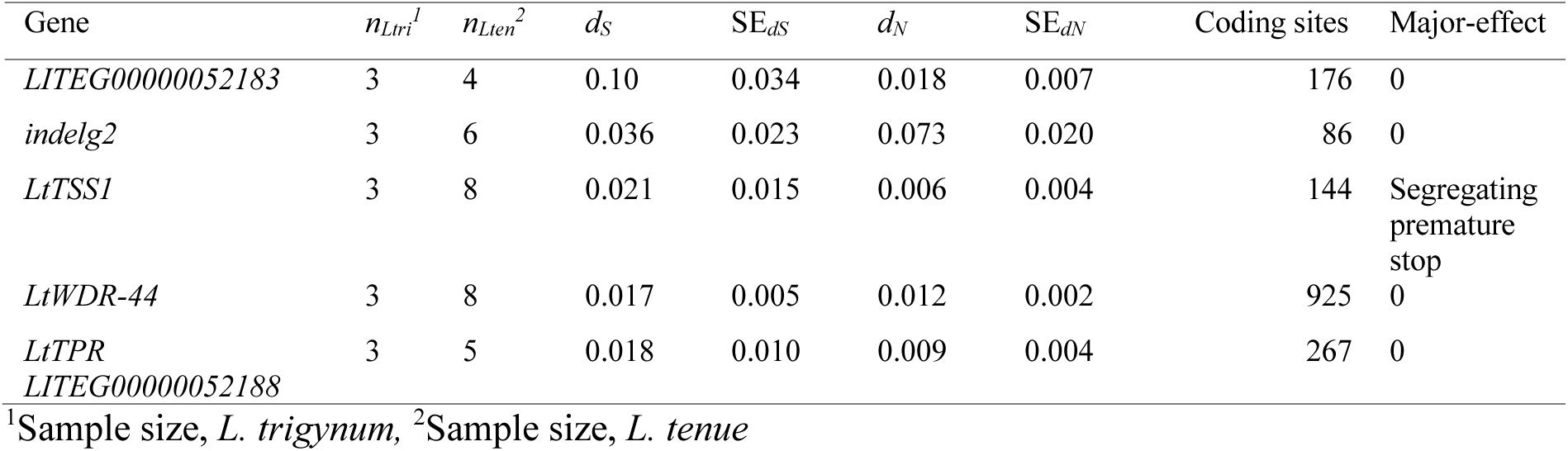
Sequence divergence between *L. trigynum* and *L. tenue* genome assemblies at the five homologous genes they share in the *S-*locus region. Synonymous and nonsynonymous substitution rates are indicated by *d_S_* and *d_N_*, respectively. Standard errors (SE) of both are given as well as the number of major-effect mutations.

### The S-linked Gene LtWDR-44 is Downregulated in Floral Buds of L. trigynum

To determine if distyly breakdown in *L. trigynum* was associated with changes in transcript abundance at candidate genes, we contrasted gene expression at *S*-linked genes in *L. tenue* thrums and *L. trigynum* homostyles. We focused on detecting altered expression of any of the genes shared between the dominant *S* allele of *L. tenue* and its derived allele fixed in *L. trigynum* (i.e, *liteg52183*, *indelg2*, *LtTSS1*, *LtWDR44* and *LtTPR/liteg52188*). Out of these genes, only the stamen length and/or male SI candidate gene *LtWDR-44* was significantly differentially expressed in floral buds, being downregulated in floral buds of *L. trigynum* compared to *L. tenue* (Log_2_-fold change = −1.58, p < 0.01, Fig. 3). *LtWDR-44* was not differentially expressed between *L. tenue* thrums and *L. trigynum* homostyles in leaves (Supplementary Fig. S4; Supplementary Information). None of the remaining *S*-linked genes showed significantly different levels of expression in floral buds or leaves between *L. tenue* thrum and homostylous *L. trigynum* (Supplementary Fig. S4; Supplementary Information).

**Fig. 3.**
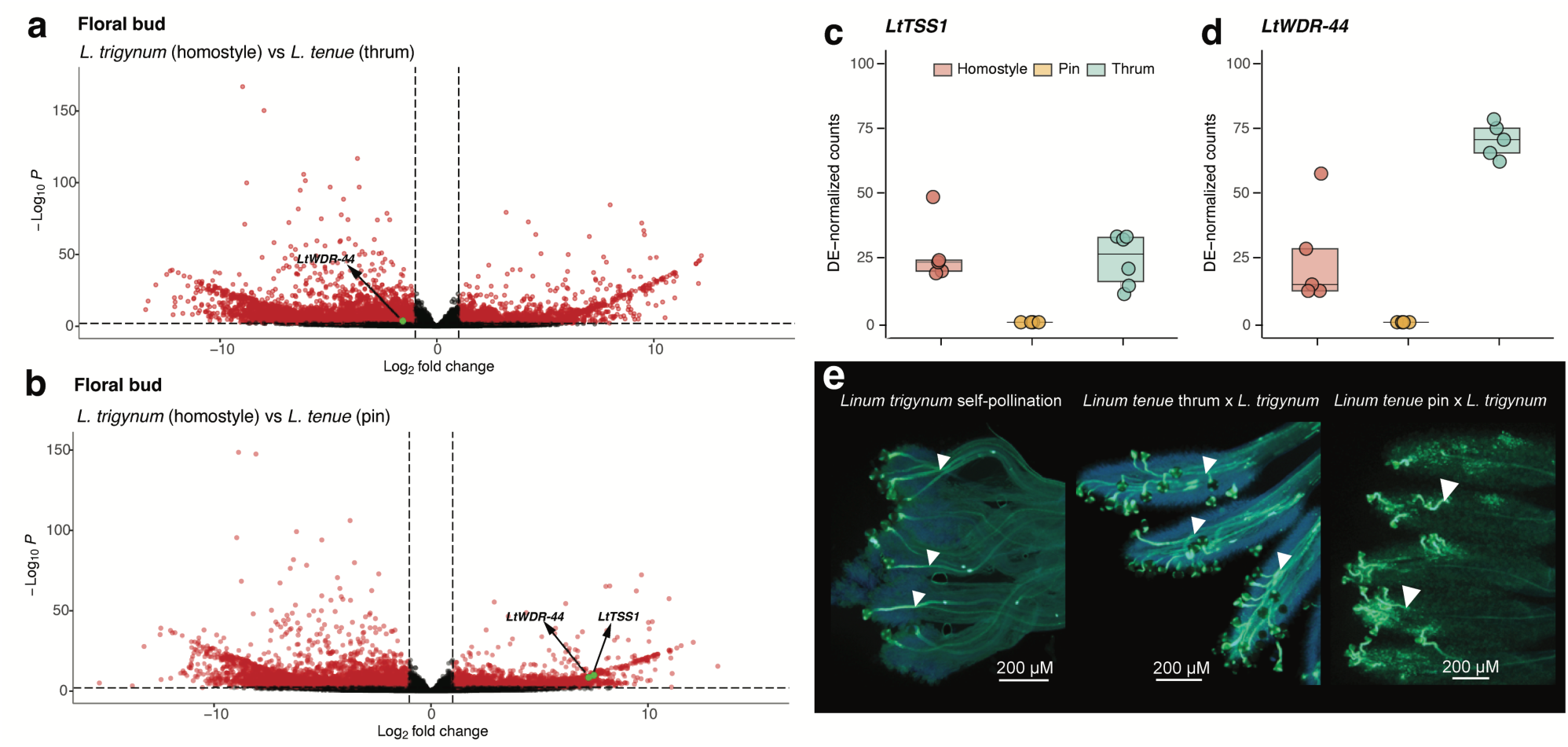
Differential expression of *S-*linked candidate genes in floral buds and pollination assays suggest a role for downregulation of *LtWDR-44* in transition to homostyly. (a) Volcano plot depicting fold change vs significance of differential expression (log_2_fold change *L. trigynum*/*L. tenue*) between *L. trigynum* homostyles and *L. tenue* thrums. Significant *S-*linked genes (only *LtWDR-44* in this analysis) are indicated. (b) Volcano plot depicting fold change vs significance of differential expression (log_2_fold change *L. trigynum*/*L. tenue*) between *L. trigynum* homostyles and *L. tenue* pins. Significant *S-*linked genes (*LtWDR-44* and *LtTSS1*) are indicated. (c) Normalized counts for *LtTSS1* in *L. trigynum* homostyles, *L. tenue* pins and *L. tenue* thrums. (d) Normalized counts for *LtWDR-44* in *L. trigynum* homostyles, *L. tenue* pins and *L. tenue* thrums. (e) Representative epifluorescence micrographs of pollination assays demonstrating self-compatibility of *L. trigynum* (left), a compatible reaction when pollinating *L. tenue* thrum (middle) but not pin (right) with *L. trigynum* pollen. Pollen tubes are indicated by white arrows. Note that the site of pollen tube rejection is in the style in *Linum*, such that an incompatible cross yields shorter, aborted pollen tubes. Scale bars indicate the degree of magnification.

### Pollination Assays Indicate Altered Male SI in *L. trigynum*

The evolution of homostyly from distyly is often associated with a shift from heteromorphic SI to SC (reviewed by Barrett 2019). If the *S-*linked gene *LtWDR-44* is a determinant of male SI, as we previously hypothesized (Gutiérrez-Valencia et al. 2022a), then we would expect downregulation of this gene to affect the male (pollen) SI expressed by *L. trigynum*. Specifically, we would expect *L. trigynum* pollen to behave similarly to *L. tenue* pin pollen and elongate successfully in thrum but not pin *L. tenue* pistils. Crossing assays followed by pollen tube staining showed that *L. trigynum* grew long pollen tubes in pistils of *L. tenue* thrum plants, but not in pistils of *L. tenue* pin plants (Fig. 3e, Supplementary Fig. S5, Supplementary Information; note that for incompatible crosses, pollen tube rejection occurs in variable sites along the style in *Linum*, but mostly in the upper part; Murray 1986). These results are consistent with expectations if downregulation of *LtWDR-44* in *L. trigynum* alters male SI function. A switch in male incompatibility phenotype from thrum to pin-type in an individual with functional thrum-type female incompatibility reaction would be expected to lead to SC.

### High Levels of Inbreeding in the Homostylous *L. trigynum*

In addition to reduced herkogamy, *L. trigynum* has other traits typical of the floral selfing syndrome, including a smaller floral display (Fig. 1) and a fivefold lower pollen:ovule ratio compared to *L. tenue* (Repplinger 2009), suggesting that self-pollination is its main mating strategy. If so, we would expect *L. trigynum* to be highly inbred compared to distylous SI *L. tenue*. To assess whether this was the case, we estimated the inbreeding coefficient using genome-wide polymorphism data from 224 individuals representing eight populations per species (average *n*=14 individuals per population; Table S1, Supplementary Information). We found that estimates of the inbreeding coefficient (*F_IS_*) for *L. trigynum* were significantly higher than those for *L. tenue* populations (Kruskal–Wallis test followed by Dunn’s test with Benjamini-Hochberg adjustment of P-values, *P* < 0.05 for all except two *L. trigynum* – *L. tenue* population comparisons; summary of median *F_IS_* values across populations: *L. tenue* mean=0.09, S.E.=0.03, *n*=8, *L. trigynum* mean=0.87, S.E.=0.04, *n*=8) (Fig. 4a). Assuming equilibrium (Wright 1969), the mean effective self-fertilization rate in *L. trigynum* is 0.93 (2 *F_IS_* /[1+ *F_IS_*]). These results show that *L. trigynum* is more inbred than *L. tenue*, consistent with expectations if SC and reduced herkogamy as a result of homostyly led to elevated selfing rates.

**Fig. 4.**
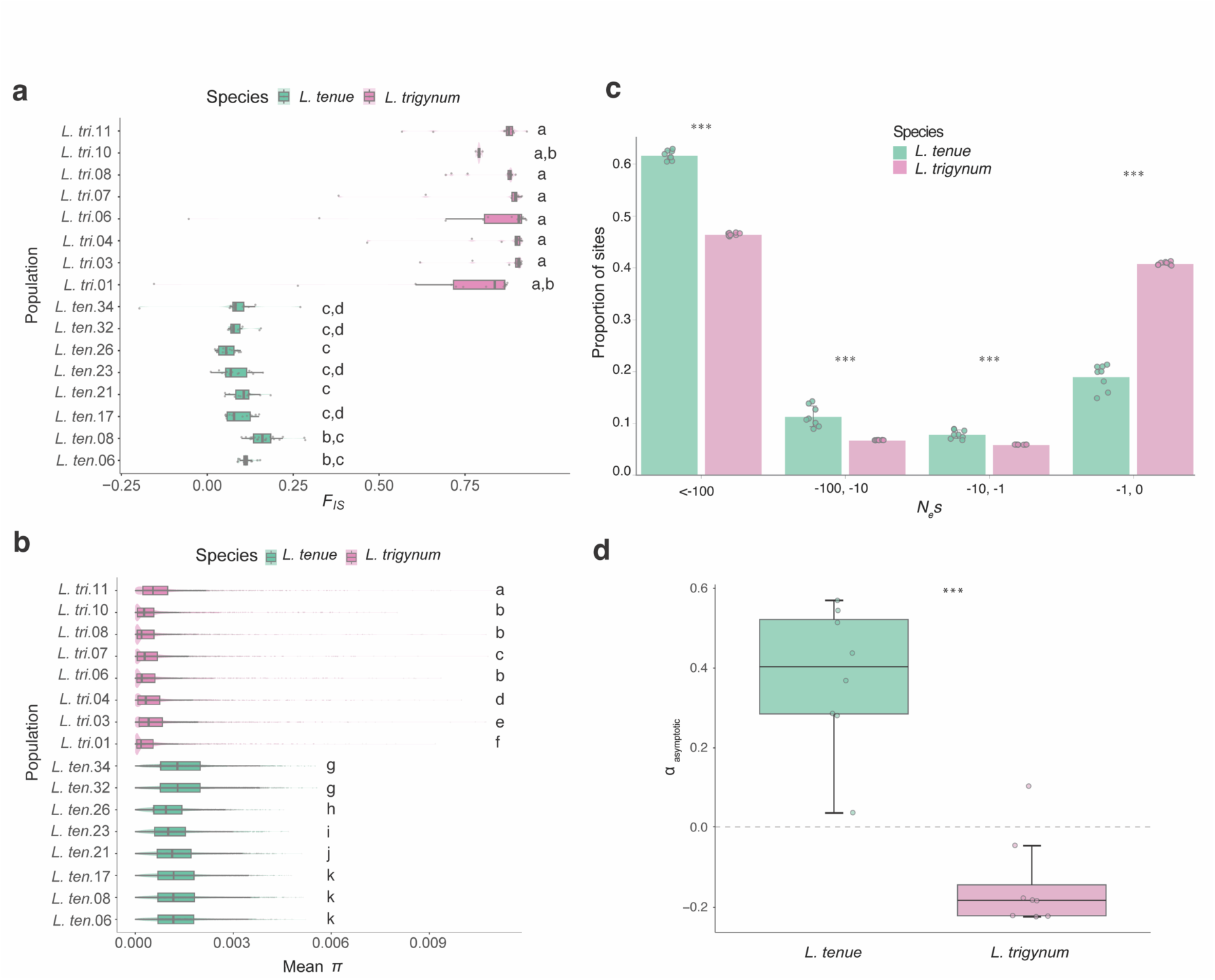
The shift to homostyly in *L. trigynum* is associated with higher levels of inbreeding, reduced genome-wide polymorphism, and a reduced efficacy of purifying and positive selection. (a) Inbreeding coefficient (*F_IS_*) estimates for *L. trigynum* and *L. tenue* indicate higher inbreeding in *L. trigynum.* Boxplots show the distribution of *F_IS_* estimates separately for each of eight *L. trigynum* populations (marked with a *L. tri.* prefix and a population number) and for each of eight *L. tenue* populations (marked with a *L. ten.* prefix and a population number, see Table S1). **(b)** Comparisons of *π* estimates in windows of 100 kb across the genome show drastically lower nucleotide diversity in *L. trigynum* compared to *L. tenue* populations. Boxplots show the distribution of *π* estimates in windows of 100 kb separately for each of eight *L. trigynum* populations (marked with a *L. tri.* prefix and a population number) and for each of eight *L. tenue* populations (marked with a *L. ten.* prefix and a population number, see Table S1). **(c)** Comparison of genome-wide distribution of negative fitness effect estimates for 0-fold degenerate nonsynonymous mutations between *L. trigynum* (*n*=8) and *L. tenue* (*n*=8). Error bars represent the standard deviation derived from populations of each species. The fraction of mutations between species was significantly different for each category of *N_e_s* (0 > *N_e_s* > −1 = effectively neutral, −1 > *N_e_s* > −10 = moderately deleterious, *N_e_s* > 10 −10 > *N_e_s* > −100 and *N_e_s* < −100 = strongly deleterious) (\*\*\**P* < 0.001, Wilcoxon rank sum test). (**d**) Comparison of alpha estimates for *L. tenue* and *L. trigynum* populations (*** *P* < 0.001, Wilcoxon rank sum test).

### Stronger Population Structure and Reduced Polymorphism in L. trigynum Compared to L. tenue

Transitions to self-fertilization are expected to result in reductions of the effective population size (*N_e_*), and thus reduced genetic diversity as well as more pronounced population structure in selfers relative to outcrossers (Wright et al. 2013). To assess if the homostylous *L. trigynum* had reduced polymorphism levels genome-wide compared to the distylous *L. tenue,* we obtained windowed estimates of nucleotide diversity (*π,* 100 kb windows) for eight populations per species. In agreement with expectation, *L. trigynum* showed markedly lower genome-wide *π* than *L. tenue* (Fig. 4b) (Kruskal–Wallis test followed by Dunn’s test with Benjamini-Hochberg adjustment of P-values, *P* < 0.001 for all *L. trigynum* – *L. tenue* populations comparisons average *π* values across populations: *L. tenue*: mean=1.3×10^-3^, S.E.=1.4×10^-4^, *n*=8, *L. trigynum*: mean=5.3×10^-4^, S.E.=1.2×10^-4^, *n*=8).

Next, we investigated population structure based on our genome-wide polymorphism data (see Fig. 5a for geographical origin of the sampled individuals). Principal component analyses (PCA), population structure analyses (when *K*=2), and TreeMix-based inference based on 76,934 SNPs in non-coding regions (see Methods for details) showed that *L. tenue* and *L. trigynum* form clearly differentiated groups (Fig. 5c; Fig. S6, Supplementary Information). Additionally, structure analyses with ADMIXTURE suggest that clustering is better defined in five groups (*K*=5): three main regional populations in *L. trigynum* that largely coincide with their geographic origin, and two *L. tenue* populations (one of them widely distributed, and a second one present in the two southernmost populations) (Fig. 5b, c).

**Fig. 5.**
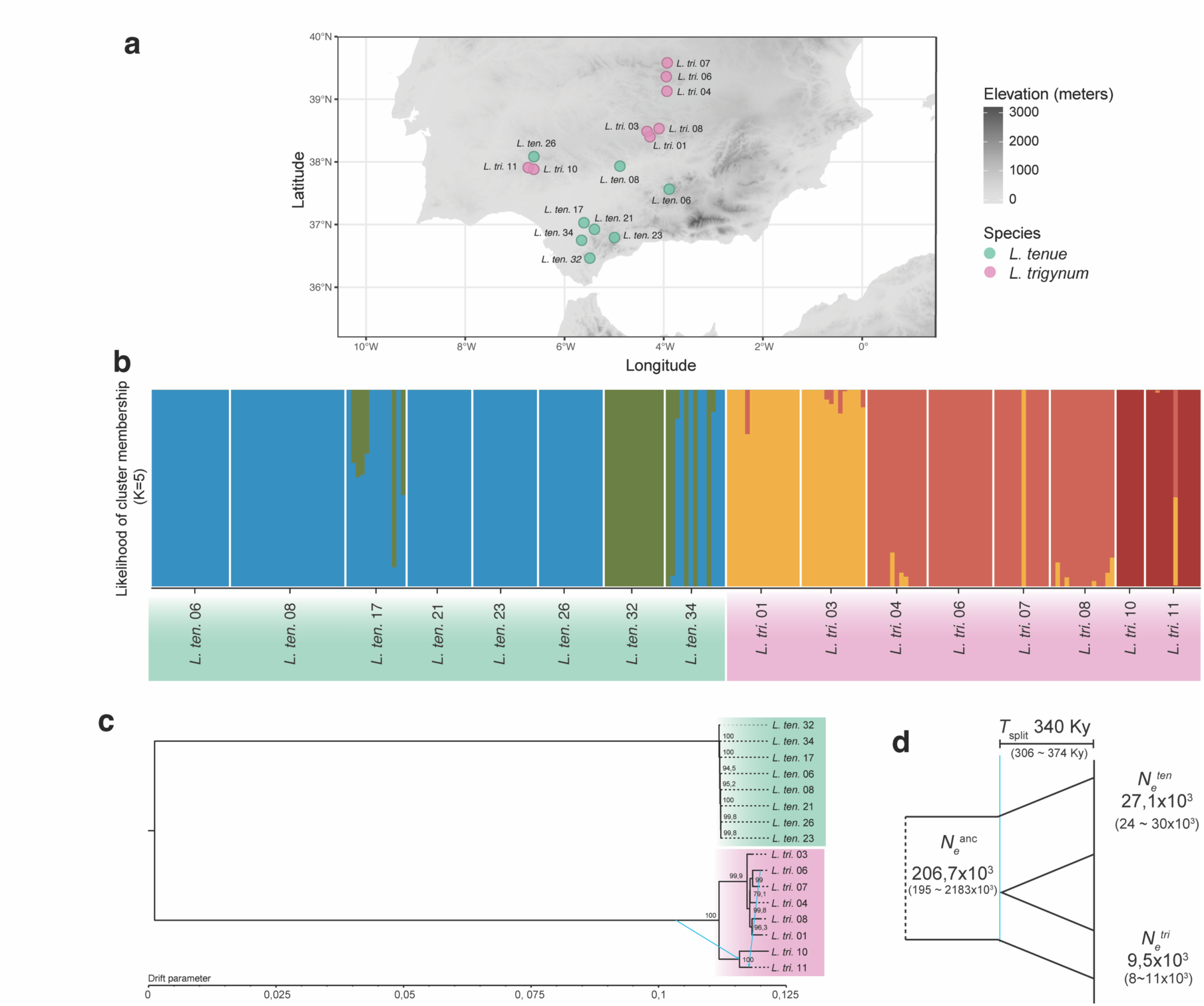
Population structure and demographic history of *L. trigynum* and *L. tenue.* **(a)** Geographic origin of the Iberian populations of *L. trigynum* (*n*=8; indicated by the prefix *L. tri.*) and *L. tenue* (*n*=8; indicated by the prefix *L. ten.*) included in this study. **(b)** Assignment of individuals to each of the five inferred ancestral clusters (*K*=5) based on admixture models to describe population structure in ADMIXTURE. **(c)** Maximum likelihood tree inferred by TreeMix. Bootstrap values for each bifurcation and two migration edges within *L. trigynum* are shown. **(d)** The demographic model inferred by *dadi*. Estimates of ancestral and current effective population sizes (in numbers of individuals) and the time of the split (in number of generations) are shown, with 95% confidence intervals based on 1000 bootstrap replicates in parentheses.

We found the highest degree of genetic differentiation between *L. tenue* and *L. trigynum* populations (*F_ST_* median=0.96, 1^st^ and 3^st^ quartile=0.95 and 0.97), followed by *L. trigynum* (*F_ST_* median=0.37, 1^st^ and 3^st^ quartile=0.28 and 0.50) and finally *L. tenue* (*F_ST_* median=0.05, 1^st^ and 3^st^ quartile=0.03 and 0.05) (Kruskal– Wallis test followed by Dunn’s test with Bonferroni corrected *P* values, *P* < 0.001) (Fig. S7, Supplementary Information). TreeMix analyses (Fig. 5c) further suggested the absence of gene flow between *L. tenue* and *L. trigynum*, and stronger population structure within *L. trigynum* than in *L. tenue*. Overall, these results show low population structure within *L. tenue*, and more marked population structure within *L. trigynum* (Fig. 5b, c).

Finally, divergence population genomic analyses in *dadi* jointly estimating demographic parameters and inbreeding levels suggested that the split between *L. tenue* and *L. trigynum* occurred about 340 kya and was associated with a marked *N_e_* reduction (Fig. 5d) and increased inbreeding in *L. trigynum* (*F_IS_* = 0.88). Together, these results suggest that the transition to homostyly was associated with higher rates of self-fertilization and reduced *N_e_,* and occurred within a relatively recent evolutionary timeframe.

### Relaxed Purifying Selection Against Weakly Deleterious Mutations in *L. trigynum*

If the shift to selfing in *L. trigynum* led to relaxed purifying selection on nonsynonymous mutations, we might expect to observe an elevated ratio of nonsynonymous to synonymous polymorphism (*π_N_/π_S_*) in *L. trigynum* relative to *L. tenue*. In line with this expectation, we found that *π_N_/π_S_* estimates were slightly higher in *L. trigynum* than in *L. tenue* (Fig. S8, Supplementary Information) (*L. tenue*: mean= 0.40, S.E.= 0.02, *n*=8, *L. trigynum*: mean= 0.50, S.E.= 0.15, *n*=8). To test if elevated *π_N_/π_S_* in *L. trigynum* was due to weaker purifying selection, we estimated the distribution of negative fitness effects (DFE) of new nonsynonymous mutations in each population of *L. trigynum* and *L. tenue* using fastDFE v1.0.0 (Fig. 4c). In line with the expectation that reduced *N_e_* in selfers should lead to relaxed selection against weakly deleterious mutations (reviewed by Wright et al. 2013), we found that the proportion of new nonsynonymous mutations that are effectively neutral was significantly higher in *L. trigynum* (0 > *N_e_s* > −1: median=0.44) compared to *L. tenue* (0 > *N_e_s* > −1: median=0.24) (Wilcoxon rank-sum test, *P* < 0.001, *n*=8 populations per species), suggesting that selection against weakly deleterious mutations is relaxed in *L. trigynum* relative to *L. tenue*. Moreover, we found a lower proportion of new nonsynonymous mutations with moderate and strongly deleterious effects in *L. trigynum* (−1 > *N_e_s* > −10: median=0.064, −10 > *N_e_s* > −100: median=0.074, *N_e_s* < −100: median=0.074) compared to *L. tenue* (−1 > *N_e_s* > −10: median=0.096, and −10 > *N_e_s* > −100: median=0.132, *N_e_s* < −100: median=0.524) (Wilcoxon rank-sum test, *P* < 0.001, *n*=8 populations per species). Together, these results show that the efficacy of purifying selection on new nonsynonymous mutations is lower in *L. trigynum* than in *L. tenue*.

### Reduced Contribution of Positive Selection to Nonsynonymous Divergence in *L. trigynum*

We compared the proportion of nonsynonymous divergence driven by positive selection (alpha) in *L. tenue* and *L. trigynum* using a method that accounts for the impact of weakly deleterious mutations on alpha (Messer and Petrov 2013). We found that estimates of alpha were significantly higher for *L. tenue* than for *L. trigynum* (Wilcoxon rank sum test, *n*=8 populations per species, *P <*0.001, median alpha: 0.40 vs −0.18, for *L. tenue* and *L. trigynum*, respectively) (Fig. 4d). For all except one *L. tenue* population, 95% confidence intervals of alpha estimates did not overlap with zero (Fig. S9, Supplementary Information), indicating that there is evidence for adaptive nonsynonymous divergence. In contrast, for *L. trigynum*, all 95% confidence intervals encompassed zero, suggesting that there is no evidence for significant adaptive nonsynonymous divergence in *L. trigynum* (Fig. S9, Supplementary Information).

## Discussion

The genetic basis and evolutionary significance of distyly has interested many generations of biologists (e.g, Darwin 1876, 1877, Bateson and Gregory 1905, Ernst 1936). The study of distyly and its breakdown has further inspired development of theory (e.g, Dowrick 1956, Charlesworth and Charlesworth 1979a, 1979b, Lloyd and Webb 1992) and vast amounts of empirical work (reviewed by Ganders1979, Barrett 1992, Barrett 2019). In recent years, new genomic tools have enabled the generation of high-quality genome assemblies of non-model plant species. This development has opened up the possibility to use genomic approaches to test long-standing hypotheses on the evolution and loss of distyly in a range of systems where such work was not possible before.

Although *Linum* is a classic system for the study of distyly (Darwin 1863), few studies have investigated the genetic causes and population genomic consequences of loss of distyly and evolution of homostyly in *Linum*. Here, we generated a high-quality reference genome assembly of the homostylous species *L. trigynum* and used it to test hypotheses on the transition to homostyly, and to investigate the population genomic effects of this shift in terms of inbreeding levels, patterns of polymorphism, and the efficacy of natural selection genome-wide.

While earlier theory posited that rare recombination events between the recessive and dominant alleles at the *S*-locus were the cause for distyly breakdown (Dowrick 1956, Charlesworth and Charlesworth 1979a), recent findings have indicated that distyly *S-*loci harbor presence-absence variation and that thrums are predominantly hemizygous at the *S*-locus rather than heterozygous (Li et al. 2016, Shore et al. 2019, Gutiérrez-Valencia et al. 2022a, Fawcett et al. 2023, Yang et al. 2023, Zhao et al. 2023). These findings make previous models of breakdown less likely and indicate that mutations either at the *S-*locus or at unlinked modifier loci are more plausible causes for the evolutionary shift from distyly to homostyly.

The presence of a genomic region in *L. trigynum* homologous to the thrum-specific dominant *S-*haplotype of *L. tenue* strongly suggests that breakdown of distyly first occurred in a thrum individual. This finding allowed us to identify candidate sequence and gene expression changes that might have contributed to the evolution of homostyly in *L. trigynum*. Although we found a premature stop codon in *LtTSS1,* which is a strong candidate gene for style length and possibly female SI in *Linum* (Gutiérrez-Valencia et al. 2022a), this mutation was not fixed in *L. trigynum*, suggesting that loss of distyly has different genetic basis in this species. The fact that *LtTSS1* alleles with and without this mutation segregate in homostylous populations suggests that it might not have altered stylar condition, possibly because it is located near the end of the coding sequence and does not disrupt the two conserved motifs present in this gene (Ushijima et al. 2012).

Our differential expression analyses showed that *LtWDR-44,* a candidate gene for anther position and/or pollen functioning, was the only *S*-linked gene downregulated in floral buds of *L. trigynum* compared to *L. tenue*. Pollination assays further showed that *L. trigynum* pollen tubes can grow in styles of *L. tenue* thrums but not in those of pins, consistent with expectations if downregulation of *LtWDR-44* in *L. trigynum* resulted in a switch in the male SI reaction from thrum-type to pin-type, a change that is likely to have rendered *L. trigynum* SC. These results suggest that mutations resulting in downregulation of the expression of *LtWDR-44* are a plausible cause of the evolution of SC and homostyly in *L. trigynum* via altered pollen SI and possibly effects on anther height, and should be subject to further functional study. Differences in TE content of the promoter region of *LtWDR-44* and especially the presence of LTR-Ty3 retrotransposons only in *L. trigynum* and not in *L. tenue* are of special interest in this respect, given that insertions of LTR-Ty3 elements in promoter regions have been associated with reduced gene expression (e.g., Steige et al. 2015, Castanera et al. 2023). Moreover, previous work showed that loss of homomorphic SI (e.g., in *Arabidopsis thaliana*) might have evolved in association with mutations in the 5’ flanking region of the male component of the *S*-locus (i.e., *SCR*) by reducing its expression in anthers (e.g., Tsuchimatsu et al. 2017). It remains to be studied if this path to SC is feasible and frequent in species with heteromorphic SI systems.

Future work could expand our understanding of the breakdown of distyly in *L. trigynum* by investigating whether the alteration of *LtWDR-44* might simultaneously alter male SI reaction and contribute to decreased herkogamy, or if more complex evolutionary pathways could have led to the evolution of homostylous flowers. Studies in *Turnera* support the plausibility of this final scenario by showing that the *S*-linked *YUCG* gene determines the functioning and size of pollen, but does not affect stamen length (Henning et al. 2022). Our finding of downregulation of *LtWDR-44* associated with SC and a switch from thrum-to pin-type male SI in *L. trigynum* constitutes an interesting contrast to the mode of loss of distyly in *Primula* and *Fagopyrum*, which involved mutations at *S-*linked candidate genes for style length and female SI function (Huu et al. 2016, Huu et al. 2022, Mora-Carrera et al. 2021, Fawcett et al. 2023).

Theory predicts that SC long homostyles should be favored during establishment of homostyly (Dowrick 1956, Charlesworth and Charlesworth 1979a), as they can spread faster than short homostyles. Work in *Primula* (Huu et al. 2016, Huu et al. 2022, Mora-Carrera et al. 2021) and *Fagopyrum* (Fawcett et al. 2023) has shown that shifts to homostyly in both lineages were most likely precipitated by mutations at *S*-linked genes that simultaneously govern pistil elongation and female SI. These results are in line with theoretical predictions (Dowrick 1956, Charlesworth and Charlesworth 1979a). In contrast, our findings in *L. trigynum* so far do not conform to these theoretical expectations, but rather suggest that mutations leading to a SC short homostyle have been fixed in this species. There are several possible reasons, which are not mutually exclusive, for why this might be the case. First, in the absence of long homostyles, short homostyles would still have an advantage over distylous individuals under conditions that favor selfing (Dowrick 1956, Charlesworth and Charlesworth 1979a), including the establishment of new populations in localities where pollination opportunities are scarce (Baker 1955, Stebbins 1957). In *Linum,* homostyly is associated with range expansion outside of the ancestral area in the Western Palearctic (Maguilla et al. 2021), suggesting that such scenarios might be plausible, including for *L. trigynum* which has a much wider geographical range than distylous *L. tenue*. If mutations resulting in homostyly are rare, as we might expect given the relatively low levels of genome-wide polymorphism in *L. trigynum* and *L. tenue*, then chance events might determine whether short or long homostyles are first formed and exposed to selection.

Second, mutational biases (reviewed in Shimizu and Tsuchimatsu 2015) might affect the rate of appearance of each of the two types of homostyles. For instance, we might expect SC long homostyles to evolve more readily if the mutation rate at genes affecting style length/female SI is higher than that at genes affecting anther height/male SI. Conversely, if the mutation rate is higher at genes affecting anther height/male SI, we could expect SC short homostyles to evolve more readily. While the length of regulatory regions affecting style length/female SI or anther height/male SI is generally not known in any system and should ideally also be considered, it is interesting to note that in *Primula vulgaris,* the gene for female SI and style length is substantially longer than the gene affecting anther position (style length/female SI gene *CYP^T^* aka *CYP734A50:* 68 kb, vs. anther height gene *GLO^T^*, aka *GLO2*: 25 kb; Li et al. 2016), whereas in *L. tenue* the situation is the opposite (*LtTSS1* gene length ∼1.9 kb, *LtWDR-44* gene length ∼11.0 kb). Thus, if mutational biases resulting from mutational target size alone are driving differences among systems in the type of homostyly that evolves, we might expect style length/female SI genes to be more frequently involved in *Primula*, and less frequently in *Linum*. To test this hypothesis, it would be beneficial to take advantage of the multiple independent transitions from distyly to homostyly that have been documented in *Linum* (at least six transitions inferred in Ruiz-Martin et al. 2018; note that in *Linum* it is not generally known *a priori* whether homostyles are short or long homostyles). In the case of *L. trigynum*, where mutations at *S*-linked coding sequences are likely not associated to distyly breakdown, differences in the length of *cis*-regulatory regions of *S*-linked genes could have similarly determined which functions of distyly were more likely to be affected via altered expression. It remains to be determined if the regulatory sequences of *LtWDR-44* are larger than those of *LtTSS1*, and if this feature could have increased the chances of regulatory mutations affecting *LtWDR-44*.

Third, it is possible that the function of *S-*locus genes in *Linum* render a transition to homostyly via long homostyles more difficult than in *Primula*. In the absence of functional genetic studies of *LtTSS1,* it remains uncertain whether loss-of-function mutations at this gene could simultaneously alter both style length and female SI, as is the case in *Primula*. A caveat regarding the potential causes of loss of distyly in *L. trigynum* that we have identified is that we cannot completely rule out a contribution of mutations at non-*S*-linked loci to reduced herkogamy and SC in *L. trigynum* (Ganders 1979, Mather and Winton 1941). Unfortunately, such studies are precluded here by the marked genome-wide differentiation between *L. trigynum* and *L. tenue* which prevents identification of genetic variants associated with floral morph using association mapping. Additionally, we were unable to obtain viable offspring from crosses of *L. trigynum* and *L. tenue*, preventing genetic mapping in interspecific F2 mapping populations. Functional assays should instead be used to validate candidate genetic changes that triggered loss of distyly in *L. trigynum*.

Transitions from distyly to homostyly are expected to frequently result in elevated self-fertilization rates, with major consequences for genetic variation and the efficacy of selection. To elucidate the timing of the split between *L. tenue* and *L. trigynum*, and the genomic consequences of evolution of homostyly, we analyzed 224 whole-genome sequences of individuals from eight populations each of *L. trigynum* and *L. tenue*. We found that homostyly was associated with predominant self-fertilization, and that the split between *L. trigynum* and *L. tenue* happened ca. 340 ky ago (∼12.5 *N_e_* generations ago, in terms of *L. tenue N_e_*), setting an upper bound for the evolution of homostyly. This estimate broadly coincides with the occurrence of major transformations in European flora attributed to the dramatic climatic oscillations that took place during the Quaternary, which resulted in range shifts for multiple plant lineages (Davis and Shaw 2001). Importantly, transitions to self-pollination have been acknowledged as adaptations that allow trailing edge populations to expand their range, especially if pollinator service is unreliable (Levin 2012), as has been inferred for other regions impacted by climate dynamics of the Quaternary (e.g., Groom et al. 2014). Indeed, *Usia* bee flies, which are relevant pollinators of various *Linum* species in the Mediterranean basin (Johnson and Dafni 1998, Ruiz-Martín et al. 2018, Pérez-Barrales and Armbruster 2023), significantly reduce their foraging capacity outside of their thermal tolerance range (21° to 31°C) (Orueta 2002). Therefore, it is likely that homostyly in *L. trigynum* was advantageous in the context of range expansion, past climatic instability and unreliable pollination services. Evidence for an association between range expansion and homostyly has been found in a recent phylogenetic study of the diversification of *Linum* (Maguilla et al. 2021).

Our population genomic analyses show that, in line with some of the most frequently reported consequences of shifts to selfing (reviewed in Cutter 2019), *L. trigynum* is more inbred, less genetically diverse, and shows stronger population structure than *L. tenue*. Reductions in *N_e_* resulting from bottlenecks and an increased impact of linked selection due to elevated inbreeding are expected to result in reduced efficacy of selection against weakly deleterious mutations in selfers (Wright et al. 2013, Burgarella and Glémin 2017, Cutter 2019). In line with this prediction, we found evidence for weaker purifying selection in *L. trigynum* than in *L. tenue* based on analyses of the distribution of fitness effects of new nonsynonymous (0-fold degenerate) mutations.

Selfing has been proposed to reduce the potential for adaptation (Stebbins 1957), and empirical population genomic studies have found evidence for slower rates of adaptive evolution in derived selfing lineages (e.g, Slotte et al. 2010, Burgarella et al. 2015). Using a version of the McDonald-Kreitman test that corrects for biases resulting from weakly deleterious mutations (Messer and Petrov 2013), we found that the proportion of nonsynonymous fixations driven by positive selection was significantly lower (and not significantly different from zero) for the homostylous *L. trigynum,* in contrast to the distylous *L. tenue* which had evidence for a significant contribution of positive selection to nonsynonymous divergence. While detailed analyses separately assessing selection on specific classes of genes, such as those involved in the evolution of the selfing syndrome or pollen competition (Gutiérrez-Valencia et al. 2022b) would be warranted for a more complete understanding of the impact of shifts to selfing on selection across the genome, our results so far demonstrate genomic signatures consistent with relaxed purifying and positive selection on nonsynonymous mutations in association with loss of distyly. These results are in line with those in a previous study on the genomic impact of loss of distyly in *Primula* (Wang et al. 2021), and with a wealth of studies on the genomic impact of shifts to selfing in other plant lineages (e.g, Slotte et al. 2010, Slotte et al. 2013, Laenen et al. 2018, Mattila et al. 2020, Yi et al. 2022). Comparative population genomic analyses of multiple shifts from distyly to homostyly in *Linum* will be valuable to better understand the general population genomic effects of this major evolutionary transition.

Our study represents the most comprehensive assessment of the genetic basis and population genomic consequences of the shift from distyly to homostyly in *Linum* so far. By generating a chromosome-scale genome assembly, we have enabled tests of long-standing hypotheses on the evolution of homostyly using genomic methods. We show that the genetic basis of homostyly is likely to be different in *L. trigynum* than in *Primula* and *Fagopyrum* and discuss potential reasons for such differences. Our results demonstrate pervasive genome-wide consequences of the shift to homostyly and elevated selfing rates, including reduced genome-wide polymorphism, stronger population structure, and a reduced efficacy of both purifying and positive selection. Taken together, this study expands the study of shifts from distyly to homostyly to a new system, provides a basis for future research on the pathways associated with loss of distyly, and broadens our knowledge on the evolutionary genomic consequences of shifts to homostyly and self-fertilization.

## Methods

### Plant Material and Sequencing

For genome sequencing and population genomic analyses, seeds and leaves of *L. trigynum* and *L. tenue* were sampled in 16 different localities in southern Spain, representing 8 populations per species (Table S1, Supplementary Information). Plants were grown in a controlled climate chamber at Stockholm University (Stockholm, Sweden) set to 16 h light at 20°C: 8 h dark at 18°C, 60% maximum humidity, 122 μE light intensity. We extracted DNA from 224 individuals for whole-genome short-read sequencing on an Illumina NovaSeq 6000 on a S4 flowcell (150-bp paired-end reads, v1.5 sequencing chemistry). For long-read and linked-read sequencing we extracted high molecular weight (HMW) DNA from young leaves using the CTAB method (as in Fulton et al. 1995), followed by two purification steps with Genomic tip 500/G (QIAGEN, Germany). To generate a high-quality genome assembly, HMW DNA of individual *Ltri*. 6-30-2 was sequenced on 2 Sequel II SMRT cells in HiFi mode, which led to the production of 27 Gb of HiFi data. This was supplemented with Dovetail Hi-C sequencing data (OmniC) for the same individual. For linked-read sequencing of two samples of *L. trigynum* (*Ltri.* 1-1-1 and *Ltri*. 1-42-2), four samples of *L. tenue* (*Lten*. CL-3-1, *Lten*. CL-75-1, *Lten*. STM-5-2, and *Lten.* STM-30-1), and one sample of the outgroup *L. maritimum* (*Lmar.* 08 thrum) we used the Chromium Genome Library preparation kit with sequencing on an Illumina HiSeqX system (paired-end 150bp read length, v2.5 sequencing chemistry). For genome annotation, we extracted total RNA from stems, leaves, floral buds, and mature flowers of individual *Ltri*. 6-30-2, and for differential expression analyses, we extracted total RNA from floral buds and leaves of *L. tenue* and *L. trigynum* (Sample size: *L. tenue* thrum=6 individuals, pin=4 individuals, *L. trigynum* homostyle=5 individuals) using the RNeasy Plant Mini Kit (QIAGEN, Germany). Libraries were sequenced on an Illumina NovaSeq S1 Sequencing System to produce paired-end 150bp read length reads. Full details on plant growth, sampling, extraction and sequencing are given in Supplementary Methods (Supplementary Material).

### De novo genome assemblies

HiFi PacBio subreads were assembled with IPA (v1.3.2) (https://github.com/PacificBiosciences/pbipa) using reads with QV20 or higher, which lead to a preliminary assembly of 103 primary contigs (total length = 498.60 Mb) and 2,280 associated contigs (total length = 100.57 Mb). To check for primary contigs that should be identified as haplotigs we used Purge Haplotigs (Roach et al. 2018). Illumina short-reads obtained from individual *Ltri*. 6-30-2 were mapped to the primary IPA assembly using minimap2 (Li 2018), and the alignments were processed with the function purge_haplotigs to generate a coverage histogram for contigs. The resulting histogram showed a unimodal distribution, suggesting that purging was not required. The assembly was scaffolded using Dovetail Hi-C data using 3D-DNA scaffolding (Dudchenko et al. 2017), which was pre-filtered to only keep contacts with a mapping quality higher than 30. The expected haploid number of chromosomes in *L. trigynum* is 10 (Pastor et al. 1990), and the resulting scaffolded assembly consisted of 13 pseudochromosomes (N50 = 47.03 Mb, Length = 498.10 Mb). This assembly was further edited to correct three misassemblies that were not supported by mapping of our PacBio reads to the *L. trigynum* assembly. The edited *L. trigynum* genome assembly was polished two times with Pilon (Walker et al. 2014) using Illumina short reads of the same individual. We screened the genome assembly for regions with high coverage and with similarity with the NCBI Plastid database (https://www.ncbi.nlm.nih.gov/genome/organelle/). We detected and hard-masked one region with plastid contamination on chromosome 6 (positions 59,960,001-60,370,000). We visualized broad-scale genome synteny between our *L. trigynum* and *L. tenue* (Gutiérrez-Valencia et al. 2022a) assemblies by aligning genome assemblies using minimap2 v2.4 (Li 2018) and plotting alignments larger than 100 kb using the R package circlize (Gu et al. 2014) (Fig. 2a).

Genomic linked-read sequences (10x Genomics) of two *L. trigynum* individuals, four *L. tenue* individuals and one *L. maritimum* individual were assembled with the Supernova pipeline (https://github.com/10XGenomics/supernova) using default parameters to obtain the output type pseudohap2. Furthermore, we included three additional *L. tenue* linked-read assemblies from Gutiérrez-Valencia et al. (2022a).

### Genome Annotation

The annotation of *L. trigynum* was made with TSEBRA (Gabriel et al. 2021) which combines the gene prediction of BRAKER1 (Hoff et al. 2016) and BRAKER2 (Brůna et al. 2021). BRAKER1 predicts genes using RNAseq data, therefore we trimmed the raw reads from four different tissues (leaves, stems, floral buds and mature flowers) with fastp (Chen et al. 2018), after we aligned them using STAR (Dobin and Gingeras 2015). BRAKER2 predict genes using protein databases, therefore we use protein data from *L. tenue* (Gutiérrez-Valencia et al. 2022a) and four additional Malpighiales species (*L. usitatissimum*, *M. esculenta*, *P. trichocarpa*, *S. purpurea*) download from phytozome (https://phytozome-next.jgi.doe.gov/) (Goodstein et al. 2012). We used AGAT (https://github.com/NBISweden/AGAT) to extract the coding DNA sequences needed to assess the genome completeness with BUSCO v5 (Manni et al. 2021). Repetitive elements were identified using RepeatMasker (Smit et al. 2013) using a custom repeat library modelled using RepeatModeler (Smit and Hubley 2008).

We carefully reannotated the *S-*locus region in *L. tenue* and the corresponding region in *L. trigynum* to improve on the automated annotation and fully consider RNA-seq evidence. This was necessary as the original annotation of genes at the *S-*locus in *L. tenue* was based on Augustus (Stanke et al. 2008) predictions without including RNA-seq evidence. Here, we therefore reannotated the *L. tenue S-*locus using RNA-seq data from the genome sequenced thrum individual generated in (Gutiérrez-Valencia et al. 2022a) and we likewise used *L. trigynum* RNA-seq evidence to annotate the corresponding region in *L. trigynum*. For this purpose, we first identified transcripts with StringTie v2.1.4 (Pertea et al. 2015) using the RNA-seq data from (Gutiérrez-Valencia et al. 2022a). Second, we predicted proteins using TransDecoder v5.7.0 (https://github.com/TransDecoder/). Third, we visually inspected the annotated genes using IGV v2.12.3 (Thorvaldsdóttir et al. 2012) retaining those well-supported by RNA-seq evidence.

In comparison to the previous *L. tenue* annotation (Gutiérrez-Valencia et al. 2022a), two genes (*indelg1* and *LtTSS1*) were modified, leaving a total of nine genes at the dominant allele of the *L. tenue S-*locus. Six genes with high similarity to repeats (*indelg3, indelg5, indelg6, indelg7, indelg8, indelg9*) that were not analyzed in Gutiérrez-Valencia et al. (2022a) were not retained in our updated annotation. In particular, our new annotation supported a one-exon gene structure of *LtTSS1* which was validated using PCR-based assays on cDNA and genomic DNA (Supplementary Methods, Supplementary Figure S3, Supplementary Information).

In the *L. trigynum S-*locus region we removed 22 genes annotated by TSEBRA because they were not well-supported by RNA-seq evidence and we manually reannotated two genes (*indelg2* and *LtTSS1*) in order to have the same intron-exon structure of the orthologous genes in *L. tenue*. Finally, we used NUCmer (Kurtz et al. 2004) to identify orthologous regions between *L. tenue* and *L. trigynum* in the *S-*locus region (Fig. 2d).

### Identification of Candidate Loss of Function *S*-Linked Mutations

Using minimap2 (Li 2018), we mapped the sequence of the *L. tenue* distyly *S*-locus (Gutiérrez-Valencia et al. 2022a) against the genome assembly of *L. trigynum* to identify and extract the sequence homologous to this region (Fig. 2b). The same approach was used to retrieve contigs containing the *S-*locus from 10X genomics supernova assemblies (two *L. trigynum* and six *L. tenue.* We conducted BLAST analyses (Camacho et al. 2009) to identify the sequences of *S*-locus genes in each assembly. Sequences of each gene were independently aligned using MUSCLE v3.8.31 (Edgar 2004), and only coding sequences were kept for further analyses. Coding sequences were aligned using codon-aware alignment in webPRANK (Löytynoja and Goldman 2010). We inspected each alignment using AliView (Larsson 2014) to identify major effect mutations (non-consensus splice sites and premature stop codons). Estimates of mean synonymous (*d_S_*) and nonsynonymous divergence (*d_N_*) between *L. tenue* and *L. trigynum* were obtained in MEGA X (Kumar et al. 2018, Stecher et al. 2020) using the Nei-Gojobori model (Nei and Gojobori 1986), with standard error estimates obtained using 1000 bootstrap replicates. Ambiguous codon positions were removed for each sequence pair.

### Sequence Processing, Mapping, Variant Calling and Filtering

Illumina short reads from 224 individuals representing populations of *L. tenue* and *L. trigynum* were quality and adaptor trimmed with *bbduk* from BBMap/BBTools (Bushnell 2015). Trimmed paired-end reads were mapped to the *L. tenue* genome assembly using BWA-MEM (v0.7.17) (Li 2013). Alignments of *L. tenue* and *L. trigynum* short reads to the *L. trigynum* reference genome were used for coverage analyses focusing on the *S-*locus region. For joint inference of demographic history and analyses of population structure, short-read sequences of both species were mapped to the *L. tenue* genome assembly (Gutiérrez-Valencia et al. 2022a). For all remaining analyses, sequences of each species were mapped to their corresponding reference genome, and processed independently in downstream analyses leading to variant calling.

Alignments with mapping quality lower than 20 were discarded, and we used *MarkDuplicates* from Picard tools v2.0.1 (Broad Institute 2019) to remove duplicated reads from the alignment. The resulting alignments were used to obtain genotype likelihoods with *mpileup*, and variants (SNPs/INDELs) and invariant sites were identified by samtools/bcftools, using the model for multiallelic and rare-variant calling (Danecek and McCarthy 2017). The VCF file was processed to keep only biallelic SNPs and invariant sites, and then filtered based on the maximum proportion of missing data (pm = 0.1) and read depth (5 < dp < 200). To avoid false heterozygous calls based on a low number of alternate alleles, we used a combination of allele balance and coverage filtering, which has previously been successful for highly repetitive plant genomes (see Laenen et al. 2018,Gutiérrez-Valencia et al. 2022b for a detailed description).

### Coverage Analyses

To investigate differences in depth of coverage between pin (n=25), thrum (n=26) (*L. tenue*) and homostyle (n=104) (*L. trigynum*) at the *S*-locus, sequences mapped to the *L. trigynum* genome assembly were processed to remove repetitive regions identified with RepeatMasker (Smit et al. 2013) using the *L. usitatissimum* repeat library. We used BEDTools (Quinlan et al. 2010) to estimate coverage for 50 kb windows across the genome. Estimates were further processed using in R to estimate normalized mean coverage across windows, and differences between morphs were tested using a Kruskal-Wallis test, followed by a post-hoc Dunn’s test with Bonferroni correction for multiple testing.

### Haplotype Network Analyses

To investigate whether loss of distyly in *L. trigynum* might have occurred repeatedly, we conducted a haplotype network analysis of two genes at the *S-*locus, *LtTSS1* and *LtWDR-44*. We used the genome annotations of *L. tenue and L. trigynum* species to extract coding sequences for these two genes. We then extracted the corresponding sequences from our short-read data using bam2consensus, a tool from the package tool bambam v.1.4 (Page et al. 2014). In total, 67 *L. tenue* and 100 *L. trigynum* individuals had sufficient coverage and were included in this analysis. We aligned all sequences using codon-aware alignment in PRANK (Löytynoja and Goldman 2010). Haplotypes and haplotype network were assessed using the R package pegas v.1.2 (Paradis 2010).

### Differential Expression Analyses

RNASeq raw reads from floral buds and leaves were processed with the function bbduk from BBMap/BBTools (Bushnell 2015) for quality and adapter trimming (parameters k=2, mink=11, ktrim=r, minlength=50, qtrim=rl, trimq=20, hdist=1, tbo, tpe). Reads were mapped and quantified with STAR (Dobin and Gingeras 2015), using the genome reference and the longest isoform per transcript from our updated annotation of *L. tenue*. Multimapping reads were discarded by using the flag outFilterMultimapNmax=1.

Files listing counts mapped to each feature (ReadsPerGene.out.tab) were further processed in R to conduct differential expression analyses using the package DESeq2 (Love et al. 2014). Two independent contrasts were conducted for floral buds and leaves separately: homostyles were first contrasted to thrums, and then to pins. We corrected for multiple testing with the Benjamini-Hochberg method, and genes with an adjusted Log2-fold change > |1.5| and p < 0.01 were considered significantly differentially expressed.

### Pollination Assays

To test the functionality of male and female SI in *L. trigynum*, we conducted controlled reciprocal crosses of *L. trigynum* (*n*=2 individuals) to both pin (*n*=2-5) and thrum (*n*=2-4) morphs of *L. tenue.* For comparison we also conducted self-pollination of *L. trigynum*, compatible (pin x thrum, thrum x pin) and incompatible pollinations (thrum x thrum, pin x pin) in *L. tenue* and negative controls without pollination. We did three technical replicates of each type of cross of two individuals. For pollination, whole pistils were removed from mature flower buds or recently opened flowers, placed on agar plates and hand pollinated. Pollen tube growth was observed after 4 h. For pollen tube staining, we adapted a protocol by (Mori et al. 2006). Specifically, hand-pollinated pistils were fixed with 9:1 ethanol: acetic acid solution 4 hours after pollination. Pistils were then hydrated with an ethanol series (70%, 50% and 30% ethanol) and softened overnight in 1 M NaOH. Pollen tube growth was observed under an UV fluorescence microscope (Olympus BX60) after staining with 0.1% (w/v) aniline blue solution in 100 mM K_3_PO_4_ and 2% glycerol.

### Population Structure and Timing of Split

The filtered VCF was pruned based on linkage disequilibrium (*r^2^*) prior to conducting structure and demographic analyses with PLINK (Chang et al. 2015) (parameters: window size in kilobase=50, variant count to shift the window at the end of each step=5, pairwise r^2^ threshold=0.5). We used the function *--pca* also implemented in PLINK (Chang et al. 2015) to conduct a Principal Component Analysis (PCA) on SNPs in the pruned VCF. The same data set was used for the structure analysis in ADMIXTURE (Alexander et al. 2009) using values of *K* ranging from 2 to 16. The most likely number of subpopulations in the population was determined after identifying the *K* value with the lowest cross-validation error. The results of both the PCA and structure analyses were plotted in R (R Core Team 2021). Weighted *F_ST_* values were calculated using the Weir and Cockerham estimator (Weir and Cockerham 1984) implemented in VCFTools using the function *--weir-fst-pop* (Danecek et al. 2011). Pairwise *F_ST_* values were then compared within and between species using a Kruskal–Wallis test followed by Dunn’s test, and *P*-values were corrected using the Bonferroni method in R (R Core Team 2021). We investigated historical relationships among populations using TreeMix (Pickrell and Pritchard 2012) with 0 to 5 migration edges, running 100 iterations to get the optimal number of migration edges. Using the evanno method implemented in the R package OptM (Evanno et al. 2005, Fitak 2021), we got an optimum of two migration edges, which were between populations of *L. trigynum*. We further ran 60 iterations with two migration edges to ensure we had the maximum likelihood tree, as well as 1000 bootstrap replicates to obtain confidence intervals. No admixture was detected between the species. Finally, we used an extension of *dadi* (Gutenkunst et al. 2009) by Blischak et al. (2020) to coestimate inbreeding and demographic parameters of a simple split model for *L. tenue* and *L. trigynum*. For simplicity, we only included one population each of *L. tenue* and *L. trigynum* (populations 32 and 11, respectively) in this analysis. We obtained 95% confidence intervals of parameter estimates in *dadi* using the Godambe Information Matrix to account for linkage, with 1000 bootstrap replicates and the eps step size parameter set to 1×10^-5^. We assumed a mutation rates of 7×10^-9^ (Ossowski et al. 2010) and a generation time of one year per generation when converting demographic parameters to units of number of individuals and years.

### Estimates of Inbreeding and Polymorphism

We estimated the inbreeding coefficient (*F_IS_*) (Wright 1951) to assist our understanding of the prevalence of selfing in *L. trigynum* using the option *--het* in VCFTools (Danecek et al. 2011). Nucleotide polymorphism (*π*) was estimated in 100 kb windows per population using pixy (Korunes and Samuk 2021). Statistical testing for significant differences in *F_IS_* and windowed *π* across populations was conducted in R (R Core Team 2021).

### Estimates of Purifying and Positive Selection

To compare the content of TEs between the genomes of *L. tenue* and *L. trigynum*, we created custom libraries of repeats using RepeatModeler (Smit and Hubley 2008). The resulting libraries were then used to identify loci harboring these repeats using RepeatMasker (Smit et al. 2013). We divided genome sequences in 50 kb windows using BEDTools (Quinlan and Hall 2010), and estimated the proportion of TEs per window in R (R Core Team 2021).

We investigated purifying selection at coding sequences using the annotation of both *L. tenue* (Gutiérrez-Valencia et al. 2022a) and *L. trigynum*. We calculated *π_N_/π_S_* using pixy with the options *--bed_file* and *--sites_file* to estimate *π* on a per-gene basis and by restricting the analyses to 0-fold and 4-fold sites. *π_N_* and *π_S_* were computed in R to obtain and compare values of *π_N_/π_S._* These sites were identified using the python script NewAnnotateRef.py (https://github.com/fabbyrob/science/tree/master/pileup_analyzers) (Williamson et al. 2014) ran separately on our annotated high-quality long-read *L. tenue* and *L. trigynum* assemblies, considering only the longest transcript per gene. Finally, estimates of DFE for each population were obtained using fastdfe (v1.0.0) (https://github.com/Sendrowski/fastDFE) (Sendrowski and Bataillon 2023), a python implementation of polyDFE (Tataru et al. 2017) which supports deleterious DFE inference from folded frequency spectra. We used the model “GammaExpParametrization” which models the DFE under a Γ distribution. In the model, we parametrized the folded SFS using the nuisance parameters with the option “get_demography” to account for demographic history effects. We ran 200 iterations to get the highest maximum likelihood fit, while confidence intervals were obtained with 300 bootstrap replicates. As before, 0-fold degenerate sites were considered to be under stronger purifying selection than 4-fold degenerate sites which were assumed to evolve neutrally. Site frequency spectra for 0- and 4-fold sites were obtained with easySFS (https://github.com/isaacovercast/easySFS) which uses a modified implementation of *dadi* (Gutenkunst et al. 2009) with the option “-a” to keep all SNPs. DFE results were summarized in four bins depicting the proportion of new mutations evolving as effectively neutral (0 > *N_e_s* > −1), moderately (−1 > *N_e_s* > −10), and strongly deleterious (−10, *N_e_s* > −100) and (−100 > *N_e_s*). As we were particularly interested in the proportion of effectively neutral mutations (0 > *N_e_s* > −1), which are not well represented in the output of the default binning script provided with fastDFE, which instead plots (0 > *4N_e_s* > −1), we rescaled the bin limits to achieve a more biologically interpretable representation of the DFE. The proportion of mutations in each category was compared between species using Wilcoxon rank-sum tests in R (R Core Team, 2021).

To compare the contribution of positive selection to nonsynonymous divergence in *L. tenue* and *L. trigynum*, we estimated the proportion of adaptive nonsynonymous divergence (alpha) using a method that accounts for and corrects for the impact of weakly deleterious nonsynonymous polymorphism (Messer and Petrov 2013). To obtain divergence differences, we first conducted a pairwise alignment of the reference genomes of *L. tenue* and *L. trigynum*, and then aligned them to a draft linked-read assembly of the distylous outgroup species *L. maritimum* (Table S3, Supplementary Information), using AnchorWave v. 1.1.1 (Song et al. 2022). We inferred ancestral states at variable sites in each focal species (*L. trigynum* or *L. tenue*) using a maximum-likelihood-based method (Keightley and Jackson 2018). Finally, we counted 4-fold and 0-fold sites, divergence differences and polymorphisms using fastDFEv1.0.0. Based on these counts, we conducted the McDonald and Kreitman test in iMKT (Murga-Moreno et al. 2019). We used Messer and Petrov’s (2013) asymptotic method to obtain point estimates and 95% confidence intervals of alpha for each *L. tenue* and *L. trigynum* population. We conducted a Wilcoxon rank sum test in R (R Core Team, 2021) to compare alpha estimates across *L. tenue* and *L. trigynum*.

## Data Availability

All sequencing data, genome assemblies and their annotation produced in this study has been uploaded to the European Nucleotide Archive (ENA) (https://www.ebi.ac.uk/ena/) under study accession number PRJEB67577.

## Acknowledgments

We thank José Ruiz-Martín for assistance with field work, Jerker Eriksson for technical assistance, Dr. Magdalena Vicens Fornés for providing samples, and Björn Nystedt for discussion of the project. This project has received funding from the European Research Council (ERC) under the European Union’s Horizon 2020 research and innovation programme (grant agreement No 757451), from the Swedish Research Council (grant no. 2019-04452) and the Erik Philip-Sörensen foundation to T.S, and from the Bergströms foundation to J.G.V, and from the Nilsson-Ehle foundation to P.W.H. Z.P. is financially supported by a Carl Tryggers fellowship (CTS 21:1471). The authors acknowledge support from the National Genomics Infrastructure (NGI) in Stockholm and Uppsala (Uppsala Genome Center, SNP&SEQ), funded by the Knut and Alice Wallenberg foundation, the Swedish Research Council and Science for Life Laboratory. We acknowledge support of the Uppsala Multidisciplinary Center for Advanced Computational Science (UPPMAX) for assistance with massively parallel sequencing and access to computational infrastructure. The computations were enabled by resources provided by the National Academic Infrastructure for Supercomputing in Sweden (NAISS), partially funded by the Swedish Research Council through grant agreement no. 2022-06725. Support by NBIS (National Bioinformatics Infrastructure Sweden) is gratefully acknowledged.

