## Supplementary Information for "Genetic causes and genomic consequences of breakdown of distyly in *Linum trigynum*"

Juanita Gutiérrez-Valencia[1], Panagiotis-Ioannis Zervakis[1], Zoé Postel[1], Marco Fracassetti[1], Aleksandra Losvik[1], Sara Mehrabi[1], Ignas Bunikis[2], Lucile Soler[3], P. William Hughes[1], Aurélie Désamoré[1], Benjamin Laenen[1], Mohamed Abdelaziz[4], Olga Vinnere Pettersson[2], Juan Arroyo[5], Tanja Slotte[1,*]

[1]Department of Ecology, Environment and Plant Sciences, Science for Life Laboratory, Stockholm University, Stockholm, Sweden
[2]Uppsala Genome Center, Department of Immunology, Genetics and Pathology, Uppsala University, Uppsala, Sweden
[3]Department of Medical Biochemistry and Microbiology, Uppsala University, National Bioinformatics Infrastructure Sweden (NBIS), Science for Life Laboratory, Uppsala University, Uppsala, Sweden
[4]Department of Genetics, University of Granada, Granada, Spain
[5]Department of Plant Biology and Ecology, University of Seville, Seville, Spain

*Corresponding author:
Tanja Slotte, Dept. of Ecology, Environment and Plant Sciences, Stockholm University, Stockholm SE-106 91 Stockholm, Sweden.

### Supplementary Methods

#### *Plant Material*

For genome sequencing and population genomic analyses, mature fruits and leaves of *L. trigynum* and *L. tenue* were sampled in 16 different localities of southern Spain, representing eight populations per species (Table S1, Supplementary Material). The material was collected in individual bags per mother plant and dried with silica gel for 24 to 48 hours. Seeds were sterilized with a wash of 10% bleach solution with liquid detergent, followed by rinsing with 70% ethanol and sterile, distilled water. Sterilized seeds were located in Murashige-Skoog medium (Sigma Aldrich, USA) and covered with a thin layer of top agar, followed by 25 days of stratification at 4°C. For germination, seeds in petri dishes were located in controlled climate chamber at Stockholm University (Stockholm, Sweden) set to 16 h light at 20°C: 8 h dark at 18°C, 60% maximum humidity, 122 μE light intensity. After eight days in the climate chamber, seedlings were transplanted to pots containing a mixture of equal parts of vermiculite and perlite for two parts of regular soil (Hasselfors Garden, Sweden). Leaves from one individual per seed family were taken with sterilized forceps, placed in 2 mL Eppendorf tubes, and flash frozen in liquid nitrogen. For expression analyses, floral buds and leaves sampled as described in Gutiérrez-Valencia et al. (2022a). Samples were used for different sequencing experiments as explained in *DNA extraction and sequencing*.

#### *DNA and RNA extraction and sequencing*

To produce whole-genome sequencing data for population genomic studies, genomic DNA from young frozen leaves was extracted using the kit Isolate II Plant DNA (Bioline, UK). DNA concentration and purity were measured using Qubit and NanoDrop respectively. Sequencing libraries were prepared from 1μg of DNA using TruSeq PCRfree DNA sample preparation kit from Illumina (cat# 20015962) targeting an insert size of 350 bp, following the manufacturers' instructions. Sequencing was conducted with a NovaSeq 6000 system from Illumina using a S4 flowcell (150-bp paired-end reads, v1.5 sequencing chemistry). Library preparation and sequencing were executed at the SNP&SEQ Technology Platform at the National Genomics Infrastructure in Uppsala (Sweden).

Three individuals of *L. trigynum* were randomly chosen to generate the data that would be further used for the production of *de novo* genome assemblies. To produce HiFi PacBio reads from sample *Ltri*. 6-30-2 (39.3606, -3.9477; Castilla de La Mancha, Spain) high Molecular Weight (HMW) DNA was extracted from young leaves using a protocol that involved a first extraction using the CTAB method (as in Fulton et al., 1995), followed by two purification steps with the kit Genomic tip 500/G (QIAGEN, Germany). DNA concentration was measured with Qubit, and DNA purity was quantified with NanoDrop. Sample integrity and fragment size distribution were examined with a pulsed-field capillary electrophoresis using a Femto Pulse system. DNA was sequenced on 2 Sequel II SMRT cells in HiFi mode, which led to the production of 27 Gb of HiFi data. Library preparation and sequencing were conducted at the Uppsala Genome Center from the National Genomics Infrastructure (NGI), Uppsala, Sweden. For sequencing with the 10X Chromium protocol for linked-reads, HMW DNA from two samples of *L. trigynum* (*Ltri.* 1-1-1 and *Ltri.* 1-42-2) from Castilla La Mancha, Spain, and four samples of *L. tenue* (*Lten*. CL-3-1, *Lten*. CL-75-1, *Lten*. STM-5-2, *Lten.* STM 30-1) from two natural populations in Castillo de Locubín, Spain (CL) and Santa María de Trassiera, Spain (STM), respectively, and one sample of *L. maritimum* was extracted using the kit Genomic-tip 100/G (QIAGEN, Germany). Sequencing libraries were prepared from 0.6 ng of DNA using the Chromium Genome Library preparation kit (cat# 120~260/58/61/62) following the manufacturers' protocol (#CG00043 Chromium Genome Reagent Kit v2 User Guide). The library was sequenced on a HiSeqX system (paired-end 150bp read length, v2.5 sequencing chemistry) at the SNP&SEQ Technology Platform at the National Genomics Infrastructure in Uppsala (Sweden).

To assist the scaffolding of the HiFi PacBio reads, we generated Dovetail Hi-C data from the same individual from which we obtained long reads (*Ltri*. 6-30-2). Young leaves were frozen with liquid nitrogen and ground. The sample was prepared for cross-linking, followed by cell lysis and proximity-ligation. The library was prepared according to manufacturer's instructions, followed by sequencing on a NovaSeq6000 system (NovaSeq Control Software 1.6.0/RTA v3.4.4, setup = 2x151 setup, workflow= NovaSeqXp, flowcell mode = S4). Sample and library preparation, and library sequencing were conducted at the Genomics Applications platform at the National Genomics Infrastructure in Stockholm, Sweden.

To annotate the genome assembly, we generated RNA sequencing (RNAseq) data from two technical replicates of four types of samples (stems, leaves, floral buds, and mature flowers) from the same individual used for long-read genome assembly. For differential expression analyses, we further extracted RNA from floral buds and leaves of *L. tenue* and *L. trigynum* (Sample size: *L. tenue* thrum=6, pin=4, *L. trigynum* homostyle=5). Total RNA was extracted using the RNEasy Plant Mini Kit (QIAGEN, Germany) following the manufacturer's instructions. We evaluated RNA quantity and quality using an Agilent Bioanalyzer 2100 (Agilent Technologies, Santa Clara, USA) with RNA Plant Nano microfluidic chips. RNAseq libraries were generated using the TruSeq stranded mRNA library preparation kit (Illumina, San Diego) including polyA selection. Libraries were sequenced on an Illumina NovaSeq S1 Sequencing System to produce paired-end 150bp read length sequences using the v1 chemistry at the SNP&SEQ Technology Platform in Uppsala (Sweden).

#### *Reannotation of LtTSS1*

We reannotated *LtTSS1* in *L. tenue* to improve on the automated annotation and fully consider RNA-seq evidence (see Methods for details). This was necessary as the original annotation was based on Augustus predictions without including RNA-seq evidence. While the exon-intron structure in our original annotation of *LtWDR-44* waas supported by RNA-seq evidence, we found that for *LtTSS1* the exon-intron structure originally predicted was not well supported by the RNA-seq data (Figure S3a-b).

We conducted additional PCR-based validation of our new annotation of *LtTSS1*. Specifically, we conducted a series of PCR reactions with primer pairs complementary to the template within the first exon in our new annotation and towards the 3'UTR (Figure S3c). As a template for the PCR reactions we used both the cDNA obtained from mRNA extracted from *L. tenue* thrum flower buds with RNeasy Plant Mini kit (Qiagen) and reverse-transcribed using SuperScript IV First-Strand Synthesis System (Invitrogen), as well as DNA extracted from *L. tenue* thrum leaves (Quick-DNA MagBead Plus kit, Zymo research). PCR was performed with MyTaq DNA Polymerase (Meridian Bioscience) using a program with initial denaturation at 95°C for 3 min, followed by 30 cycles of 95°C for 2s, 55°C for 2s and 72°C for 3s, and final extension at 72°C for 1 min. Reaction products were separated on 2% agarose gel and stained with GelRed (Biotium).

We found that, in contrast to expectations given the exon-intron structure in the original annotation, PCR products amplified from cDNA and genomic DNA were the same length for all primer pairs located before primer R6 (Figure S3c-d), supporting our new annotation of *LtTSS1*.

#### *References*

Fulton, T. M., Chunwongse, J., & Tanksley, S. D. (1995). Microprep protocol for extraction of DNA from tomato and other herbaceous plants. *Plant Molecular Biology Reporter*, *13*, 207–209.

Gutiérrez-Valencia, J., Fracassetti, M., Berdan, E. L., Bunikis, I., Soler, L., Dainat, J., Kutschera, V. E., Losvik, A., Désamoré, A., Hughes, P. W., Foroozani, A., Laenen, B., Pesquet, E., Abdelaziz, M., Pettersson, O. V., Nystedt, B., Brennan, A. C., Arroyo, J., & Slotte, T. (2022a). Genomic analyses of the *Linum* distyly supergene reveal convergent evolution at the molecular level. *Current Biology*, *32*, 4360-4371.

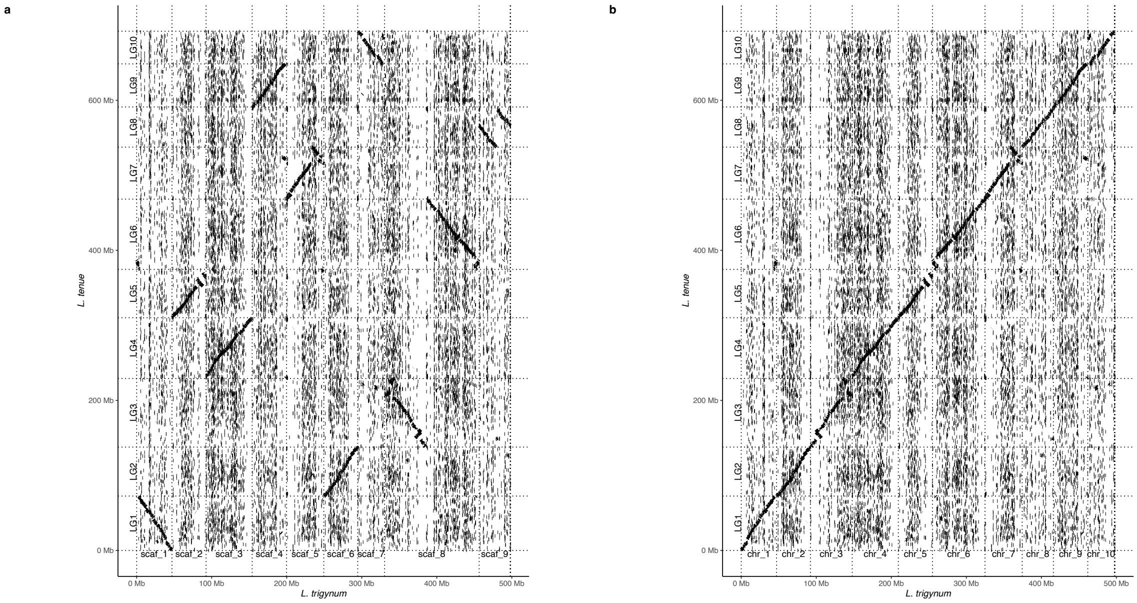


**Fig. S1.** Synteny between *L. tenue* and *L. trigynum* assemblies. (a) Alignment between the *L. tenue* genome assembly and the scaffolded *L. trigynum* assembly (b) Alignment between the *L. tenue* genome and the final *L. trigynum* assembly. Both the alignments were produced using minimap2 v2.4, and visualized with pafr (https://github.com/dwinter/pafr).

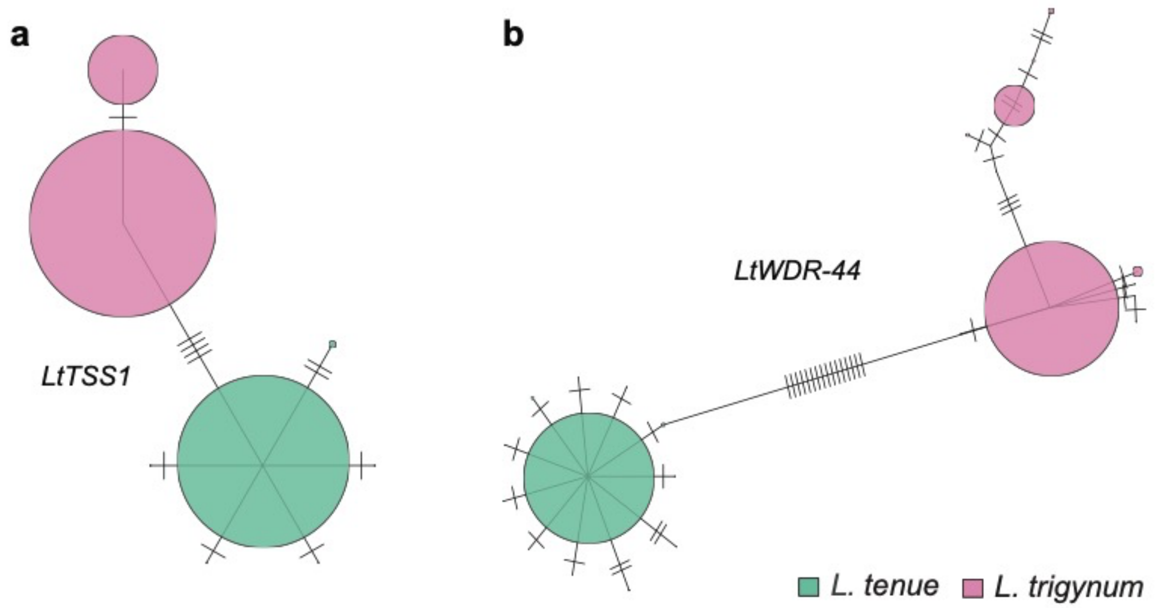


**Fig. S2.** Haplotype networks for *LtTSS1* and *LtWDR-44* based on population genomic data. (a) Haplotype network of *LtTSS1*. (b) Haplotype network of *LtWDR-44*. For both genes, the *L. trigynum S*-locus haplotype is connected to only one haplotype of *L. tenue*, consistent with a single *S*-haplotype from the distylous ancestor contributing to genetic variation at the *L. trigynum S*-locus. The size of each circle is proportional to the number of chromosomes harboring each haplotype.

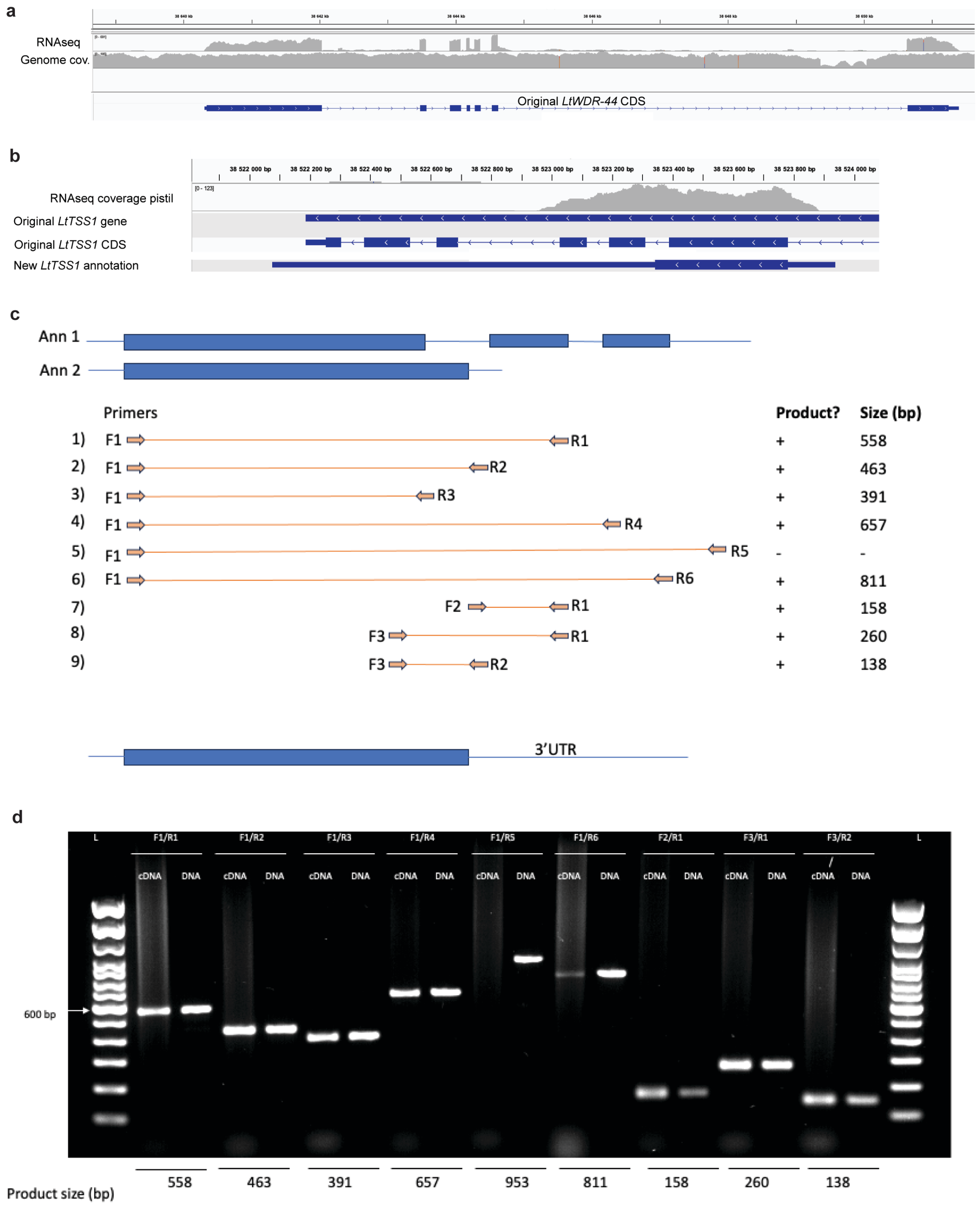


**Fig. S3.** Reannotation and validation. (a) The original annotation of *LtWDR-44* was well supported by RNA-seq data. (b) The original annotation (annotation 1) of *LtTSS1* was not well supported by RNA-seq coverage, as coverage only extends to the first three exons, and the exon-intron structure does not seem supported. (b) Locations of PCR primers in relation to exons 1-3 of the original annotation (Ann 1) and exon 1 of our new annotation of *LtTSS1* (Ann 2). The supported gene model of *LtTSS1* is shown at the bottom. (c) PCR results of selected primer pairs using as a template cDNA synthesized based on mRNA from flower buds and DNA extracted from leaves of *L. tenue* thrum individuals. DNA ladder (L) = 100bp DNA ladder, Thermo Fisher Scientific. These assays support our new annotation of *LtTSS1* with one exon, and delimit the length of the 3'UTR (based on reaction 5 which yielded no amplification from a cDNA template in contrast to amplification from genomic DNA which was successful).

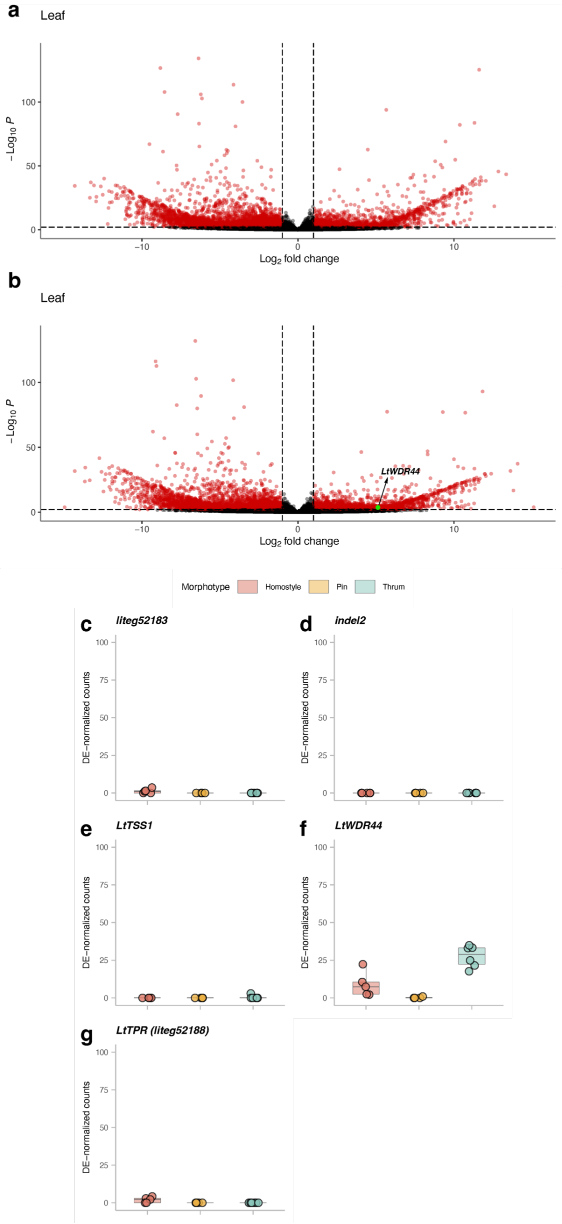


**Fig. S4. Differential expression of *S*-linked genes in leaves.** (a) Volcano plot showing fold change vs significance of differential expression between *L. trigynum* (homostyle) and *L. tenue* thrum. (b) Volcano plot showing fold change vs significance of differential expression between *L. trigynum* (homostyle) and *L. tenue* pin. Candidate genes with significant differential expression are marked in the plot. (c-e) Expression levels of *S*-linked candidate genes in *L. trigynum*, *L. tenue* pin and thrum individuals.

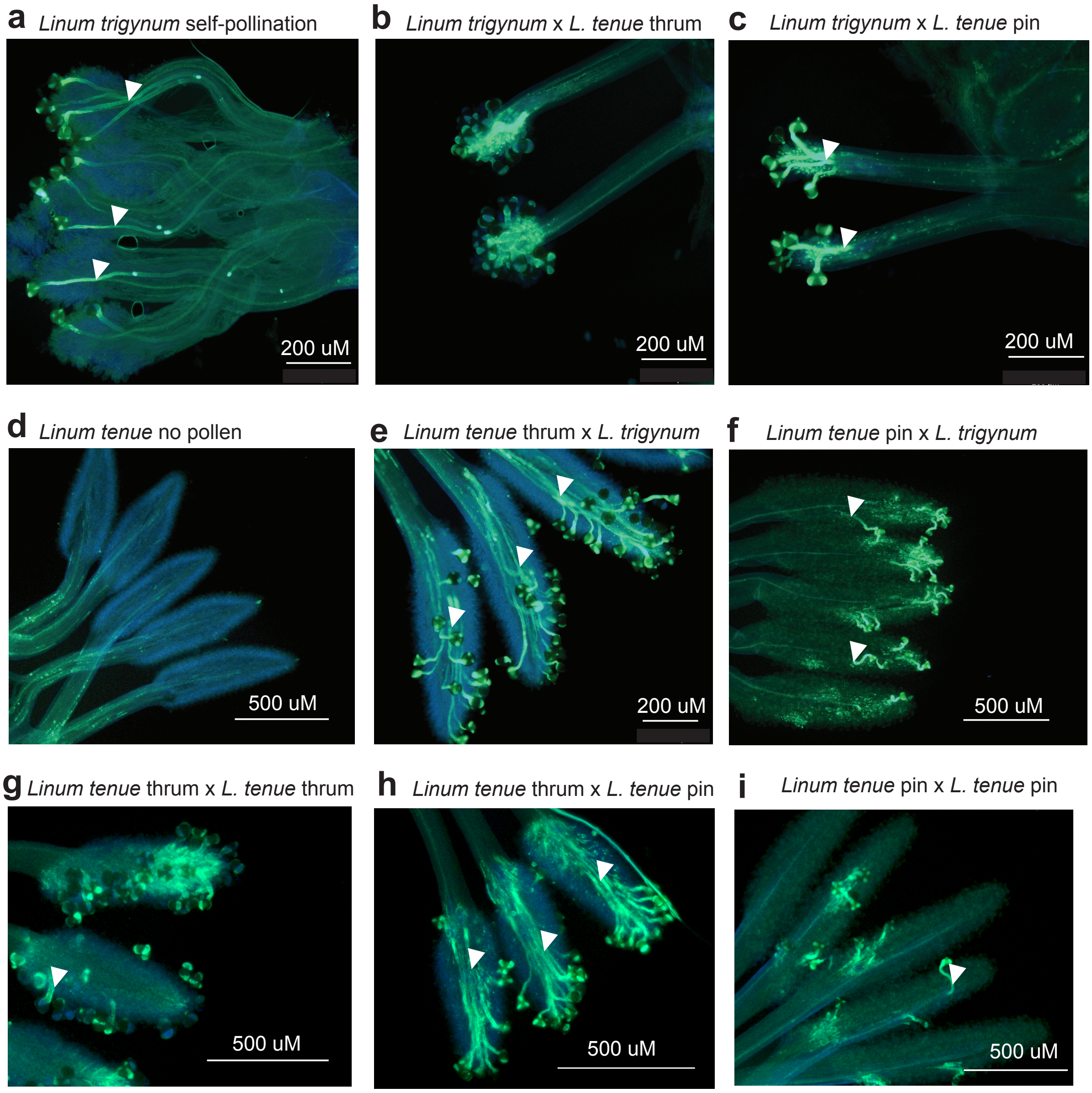


**Fig. S5.** Pollination assays indicate altered male SI function in *L. trigynum*. Fluorescence micrographs depicting pollen tube growth in styles after controlled crosses. Panels a) and d) show positive and negative controls, respectively. Panels b) and c) show weak or no pollen tube growth of *L. tenue* pollen on *L. trigynum* stigmas. Panels e) and f) show that *L. trigynum* pollen has full pollen tube growth on *L. tenue* thrum but not pin stigmas. Panels g) and i) depict incompatible *L. tenue* crosses, and panel h) a compatible *L. tenue* cross. Pollen tube growth of *L. trigynum* pollen on *L. tenue* thrum stigmas (e) resembles the outcome of a compatible *L. tenue* x *L. trigynum* cross (h). Scale bars indicate the degree of magnification, and arrows indicate pollen tubes. Note that the site of rejection in incompatible *Linum* crosses is in the style.

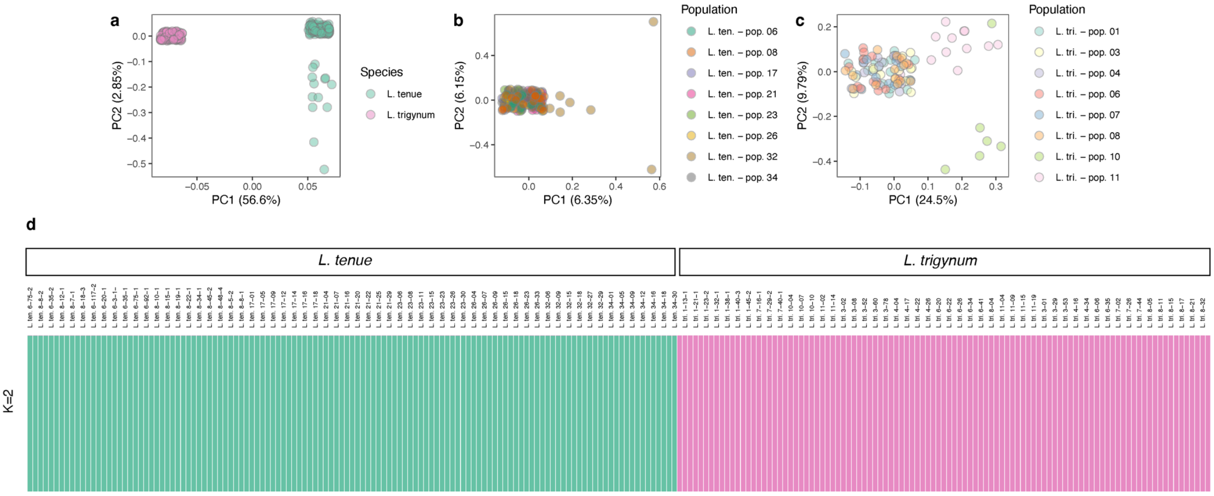


**Fig. S6.** Principal component analyses of genome-wide non-coding SNPs. Principal component analyses of 76,934 SNPs indicate that (a) *L. tenue* and *L. trigynum* are clearly separated along the first principal component (explaining 56.6% of variation), and populations of *L. tenue* cluster more closely together (b) than those of *L. trigynum* (c), consistent with weaker population structure in *L. tenue* than *L. trigynum*. (d) When only *K*=2 are considered, population structure analyses with ADMIXTURE place all *L. tenue* individuals and all *L. trigynum* individuals in two separate clusters.

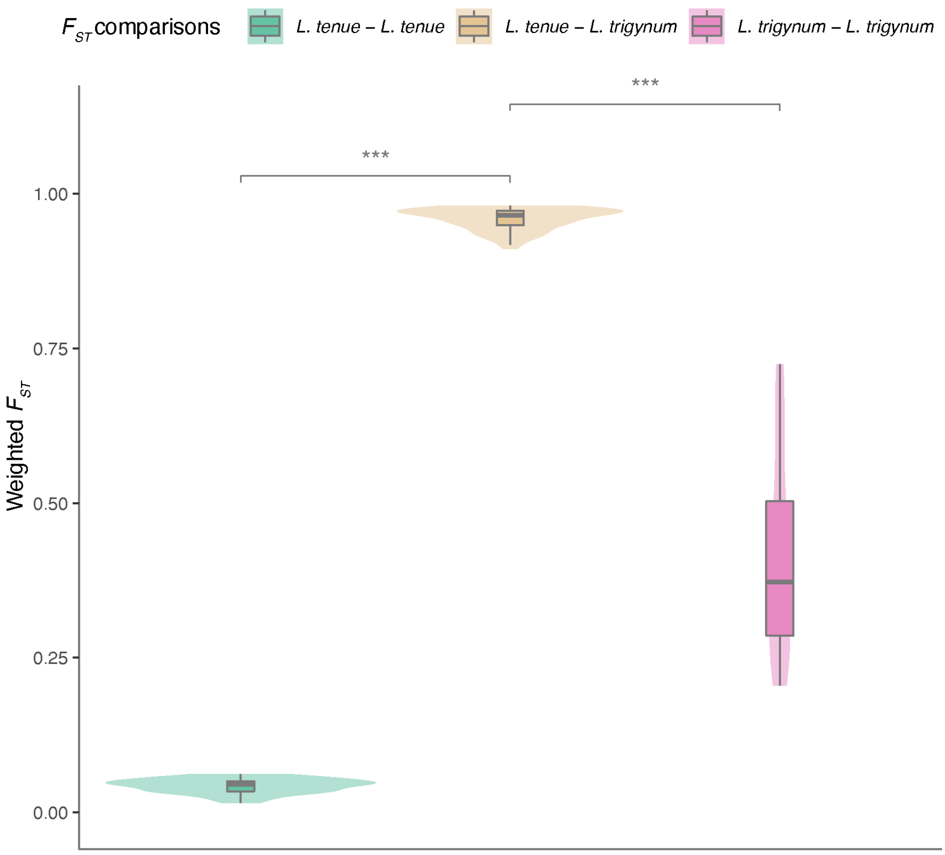


**Fig. S7.** Comparisons of $F_{ST}$ estimates within *L. tenue* and *L. trigynum*, and between populations of both species. $F_{ST}$ estimates between *L. tenue* and *L. trigynum* populations were the highest (median=0.96, 1st and 3st quantiles=0.95 and 0.97), followed by estimates within *L. trigynum* (median=0.37, 1st and 3st quantile=0.28 and 0.50), and the lowest $F_{ST}$ values were obtained for *L. tenue* populations (median=0.05, 1st and 3st quantile=0.03 and 0.05). ***$P < 0.001$ Kruskal–Wallis test followed by Dunn's test after Bonferroni correction.

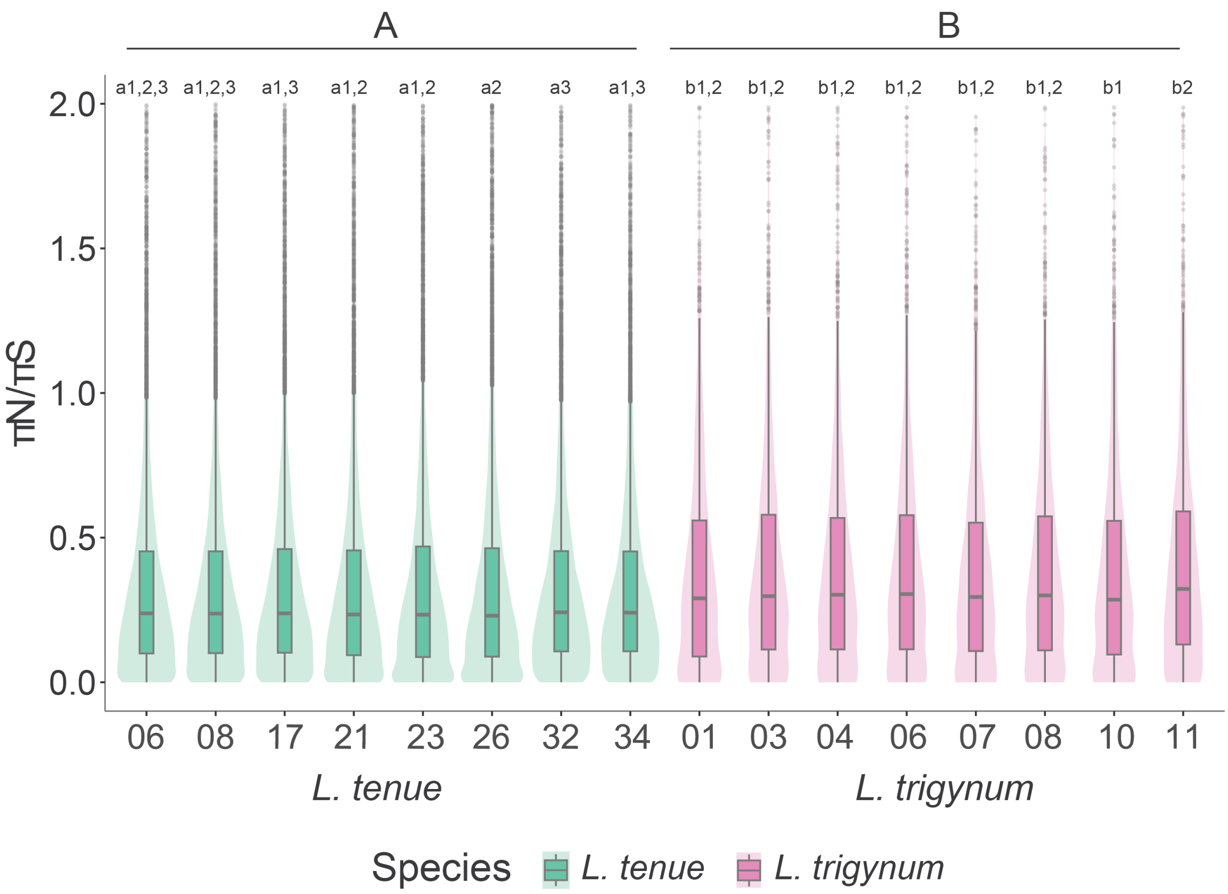


**Fig. S8.** Distribution of ratios of nonsynonymous to synonymous polymorphism in populations of *L. tenue* and *L. trigynum*. Different letters denote statistically significant $\pi_N$ / $\pi_S$ estimates between populations and between species; Kruskal–Wallis test followed by Dunn's test with Benjamini-Hochberg corrected *P* values, $P < 0.05$. $\pi_N/\pi_S$ estimates for *L. tenue* and *L. trigynum* populations. Comparisons of $\pi_N$ / $\pi_S$ estimates show that *L. tenue* (A, mean=0.40, S.E.=0.02, n=8) exhibits a significantly higher ratio of nonsynonymous to synonymous nucleotide polymorphism compared to *L. trigynum* (B, mean=0.50, S.E.=0.15, n=8).

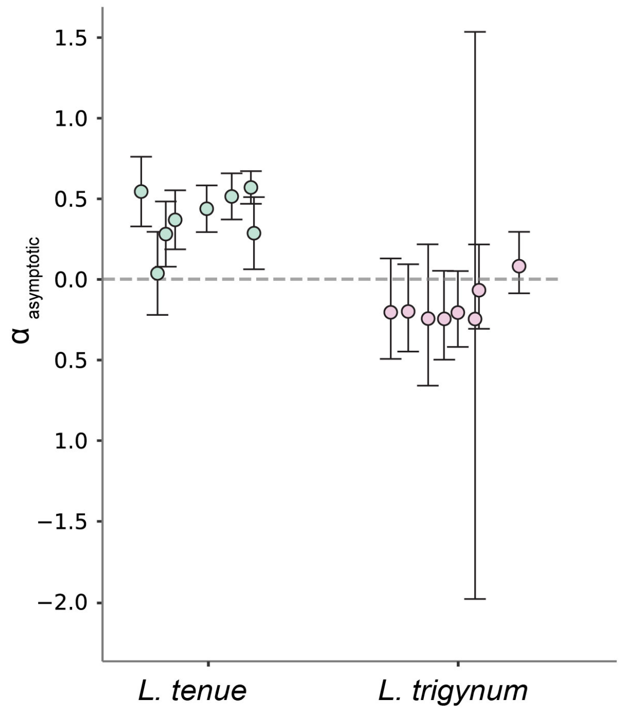


**Fig. S9.** Comparison of the contribution of positive selection to nonsynonymous divergence in *L. tenue* and *L. trigynum*. The plot shows point estimates of alpha, estimated using Messer and Petrov's (2013) asymptotic method, along with 95% confidence intervals (CIs) estimated assuming a Poisson distribution of deleterious fixations. Note that estimation of alpha in population *L. trigynum Ltri.*10 (with the widest CI) triggered a warning due to low numbers of polymorphic sites.

**Table S1.** List of populations. Populations represented in the data set used to investigate the genomic consequences of mating system shift in *L. trigynum*. The number of individuals sequenced is indicated by n. The Source column gives the study accession number at the European Nucleotide Archive (https://www.ebi.ac.uk/ena/).

| Species and Population | Population identifier | n | Latitude | Longitude | Locality | Province | Country | Source |
|---|---|---|---|---|---|---|---|---|
| *L. tenue* - pop. 17 | *L. ten.* 17 | 14 | 37.0286 | -5.6084 | El Coronil, Sevilla | Andalusia | Spain | ENA: PRJEB67577 |
| *L. tenue* - pop. 21 | *L. ten.* 21 | 14 | 36.9238 | -5.3960 | La Muela, Cádiz | Andalusia | Spain | ENA: PRJEB67577 |
| *L. tenue* - pop. 23 | *L. ten.* 23 | 14 | 36.7923 | -4.9920 | El Burgo, Málaga | Andalusia | Spain | ENA: PRJEB67577 |
| *L. tenue* - pop. 26 | *L. ten.* 26 | 14 | 37.9053 | -6.6166 | Los Marines, Huelva | Andalusia | Spain | ENA: PRJEB67577 |
| *L. tenue* - pop. 32 | *L. ten.* 32 | 13 | 36.4637 | -5.4898 | Jimena de la Frontera, Cádiz | Andalusia | Spain | ENA: PRJEB67577 |
| *L. tenue* - pop. 34 | *L. ten.* 34 | 13 | 36.7486 | -5.6552 | Arcos de la Frontera, Cádiz | Andalusia | Spain | ENA: PRJEB67577 |
| *L. tenue* - pop. 06 | *L. ten.* 06 | 26 | 37.5640 | -3.8897 | Castillo de Locubín, Valdepeñas de Jaén, Jaén | Andalusia | Spain | ENA: PRJEB52918 |
| *L. tenue* - pop. 08 | *L. ten.* 08 | 17 | 37.9320 | -4.8833 | Santa María de Trassierra, Sierra Morena | Córdoba | Spain | ENA: PRJEB52918 |
| *L. trigynum* - pop. 01 | *L. tri.* 01 | 16 | 38.4008 | -4.2775 | Fuencaliente, Ciudad Real | Castilla de La Mancha | Spain | ENA: PRJEB67577 |
| *L. trigynum* - pop. 10 | *L. tri.* 10 | 6 | 37.8816 | -6.6190 | Linares de la Sierra, Huelva | Andalusia | Spain | ENA: PRJEB67577 |
| *L. trigynum* - pop. 11 | *L. tri.* 11 | 12 | 37.9090 | -6.7314 | Jabugo, Huelva | Andalusia | Spain | ENA: PRJEB67577 |
| *L. trigynum* - pop. 03 | *L. tri.* 03 | 14 | 38.4864 | -4.3357 | Cam. de Ventillas, Fuencaliente, Ciudad Real | Castilla de La Mancha | Spain | ENA: PRJEB67577 |
| *L. trigynum* - pop. 04 | *L. tri.* 04 | 13 | 39.1280 | -3.9347 | Fernán Caballero, Ciudad Real | Castilla de La Mancha | Spain | ENA: PRJEB67577 |
| *L. trigynum* - pop. 06 | *L. tri.* 06 | 14 | 39.3606 | -3.9477 | Los Yébenes, Toledo | Castilla de La Mancha | Spain | ENA: PRJEB67577 |
| *L. trigynum* - pop. 07 | *L. tri.* 07 | 12 | 39.5816 | -3.9287 | Marjaliza, Toledo | Castilla de La Mancha | Spain | ENA: PRJEB67577 |
| *L. trigynum* - pop. 08 | *L. tri.* 08 | 14 | 38.5314 | -4.0951 | Mestanza, Ciudad Real | Castilla de La Mancha | Spain | ENA: PRJEB67577 |

**Table S2.** Assembly statistics for the *L. trigynum* genome assembly. These statistics concern the assembly based on PacBio HiFi and OmniC data.

| Assembly step | Primary assembly (Mbp) | No. sequences | N50 (Mbp) | L50 | BUSCO score (eukaryota_odb10) |
|---|---|---|---|---|---|
| IPA | 498.1 | 103 | 12.6 | 12 | C: 96.0% [S: 66.0%, D:30%], F: 0.9%, M:3.0%. |
| IPA + 3D-DNA scaffolding with OmniC data | 498.1 | 13 | 47.0 | 4 | C:95.6%[S:66.2%,D:29.4%],F:1.2%,M:3.2% |
| Final assembly | 498.1 | 15 | 47.0 | 5 | C:95.6%[S:66.2%,D:29.4%],F:1.2%,M:3.2% |

**Table S3.** Assembly statistics for Supernova assemblies. These statistics concern 10x linked-read read draft assemblies.

| **Name**[1] | **ENA**[2] | **p10**[3] | **m10**[4] | **scaffold_N50**[5] | **L50_tot**[6] | **min_scaf_L50_tot**[7] | **effective_coverage_median**[8] | **assembly_size**[9] | **est_genome_size**[10] | **Hetdist**[11] |
|---|---|---|---|---|---|---|---|---|---|---|
| Lten_CL-3-1 | this study | 59.8529 | 15.8589 | 122358 | 819 | 130000 | 32.4453 | 522480488 | 669217000 | 231 |
| Lten_CL-75-1 | this study | 130.5 | 15.5328 | 173115 | 543 | 190000 | 32.9326 | 531458269 | 650952000 | 247 |
| Lten_STM-5-2 | this study | 60.2059 | 15.5481 | 111376 | 975 | 120000 | 37.2382 | 534978380 | 639943000 | 300 |
| Lten_STM-18-3 | GCA_946150905, GCA_946150875 | 178.088 | 16.7389 | 157491 | 557 | 170000 | 28.0475 | 514042692 | 643490000 | 228 |
| Lten_STM-21-2 | GCA_946151035.1, GCA_946151015.1 | 42.8235 | 23.5384 | 80702 | 1073 | 90000 | 24.1663 | 437801162 | 650493000 | 212 |
| Lten_STM-30-1 | this study | 127.235 | 16.5916 | 268883 | 245 | 3,00E+05 | 36.2522 | 473121657 | 647544000 | 223 |
| Lten_STM-50-1 | GCA_946150985, GCA_946151025 | 37.9412 | 17.7777 | 123569 | 758 | 140000 | 29.1046 | 503721094 | 645600000 | 217 |
| Ltri_1-1-1 | this study | 78.2647 | 3.19013 | 4821513 | 33 | 4860000 | 49.5904 | 461688410 | 653639000 | 11164 |
| Ltri_1-42-2 | this study | 77.4706 | 3.66056 | 4468862 | 32 | 4460000 | 60.7739 | 463460708 | 648321000 | 8009 |
| Lmar-08 | this study | 15.3235 | 35.0141 | 33037 | 5977 | 29503 | 13.4518 | 601440318 | 1288340000 | 4575 |

[1]Name used in this study.
[2]ENA accession numbers of the two pseudohaplotypes.
**[3]**p10: for an average point on the genome, the estimated number of molecules that extend 10kb in both directions from that point, counting both alleles.
[4]m10: the estimated percent of genomic kmers that are either missing from the assembly entirely or present only in scaffolds shorter than 10 kb.
[5]scaffold_N50: the N50 size of the compressed scaffolds.
[6]L50_tot: number of scaffolds containing half of the genome length.
[7]min_scaf_L50_tot: length of the smallest L50 scaffold.
[8]effective_coverage: estimated effective coverage. This is the estimated deduplicated coverage of an average base on the genome, counting both alleles.
[9]assembly_size: the number of bases in the compressed scaffolds.
[10]est_genome_size: estimated genome size in bases, computed from the distribution of kmers.
[11]hetdist: mean distance in bases between heterozygous sites.

**Table S4.** Repeat annotation in the promoter region 2 kb upstream of the transcription start site of the two *S*-locus genes *LtWDR-44* and *LtTSS1*on pseudochromosome 10 in *L. tenue* and *L. trigynum*.

| Gene | Species | Start position | End position | Classification |
|---|---|---|---|---|
| *LtWDR-44* | *L. trigynum* | 30612087 | 30612173 | LTR/Ty3 |
| | | 30612113 | 30612174 | Unknown |
| | | 30612602 | 30613000 | LTR/Ty3 |
| | | 30613084 | 30613226 | Unknown |
| | | 30613262 | 30613418 | Unknown |
| | | 30613503 | 30613564 | DNA/hAT-Tip100 |
| | | 30613568 | 30613722 | LINE/L1 |
| | *L. tenue* | 38638631 | 38638778 | Unknown |
| | | 38638779 | 38639068 | DNA/MULE-MuDR |
| | | 38639564 | 38639857 | DNA/MULE-MuDR |
| | | 38639844 | 38640021 | Unknown |
| | | 38640008 | 38640176 | LINE/L1 |
| *LtTSS1* | *L. trigynum* | 30534704 | 30534759 | LTR |
| | | 30534776 | 30535157 | LTR |
| | | 30535398 | 30536104 | DNA/hAT-Ac |
| | | 30536271 | 30538731 | LTR/Copia |
| | *L. tenue* | 38523985 | 38524034 | LTR |
| | | 38524015 | 38524084 | LTR |
| | | 38524100 | 38524139 | LTR |
| | | 38524109 | 38524467 | LTR/Copia |
| | | 38524693 | 38525413 | DNA/hAT-Ac |
| | | 38525574 | 38526300 | LTR/Copia |
